# Peripheries in alpine phylogeography

**DOI:** 10.1101/2025.09.15.676213

**Authors:** Stéphanie Morelon, Philippe Juillerat, Julia Bilat, Hélène Mottaz, Thomas Bulliard, Florian Boucher, Laurent Juillerat, Jason Grant, Sergio Rasmann, Jeremy Gauthier, Nadir Alvarez

## Abstract

**Aim:** Peripheral regions of the Alps are often overlooked in molecular studies, yet they may play a major role in shaping the current distribution of species and genetic lineages.

**Location:** Europe

**Taxon:** Angiosperms (Primulaceae: Primula)

**Methods:** By focusing on the bear’s ear (*Primula auricula*) complex as model species, we used genetic inferences for population genetic structure and performed genetic reconstructions, species delimitations and divergence time estimates in order to get a detailed view of its molecular evolution and current genetic structuring across the Alps *sensu lato*.

**Results:** The Lombardian Alps and the southern eastern Alps are genetically distinct in the vicinity of the Adige valley. The northern and western Alps and their peripheries constitute a third clade and are separated by the siliceous central Alps. Within the latter, an additional cluster made of singular populations from the Dévoluy and Vercors regions is retrieved, likely reflecting a strong founder effect rather than an ancient divergence.

**Main conclusions:** The biogeographic history *P. auricula sensu lato* pinpoints the importance of the peripheral regions in a phylogeographic context. Populations from northern peripheral regions exhibit long-lasting isolation and *in situ* survival during glaciations, followed by recolonization into the central Alpine massif.

## Introduction

Throughout recent Earth’s history, climate shifts have profoundly shaped the biogeography of species, forcing them to migrate, adapt or face extinction as their habitat transforms (Nogués-Bravo et al., 2018).

For instance, several examples in the literature have shown that during important climate change events such as those experienced by most organisms across Pleistocene climatic oscillations, alpine plants embark on complex migration roads, shaped by both geographic and climatic barriers (Smyčka et al., 2017). These migration processes are not uniform; they vary significantly depending on the species’ ecological requirements, dispersal abilities, and their migratory starting point (Carnicero et al., 2022; Smyčka et al., 2022). Successive migrations in isolated areas and interplays might create unique diversification patterns and complex histories in mountain plants including complete and incomplete allopatric differentiation (Španiel et al., 2012; Schneeweiss et al., 2017; Rogivue et al., 2018; Boucher et al., 2021; Smyčka et al., 2022; Voisin et al., 2023). These are influenced by a combination of mechanisms such as genetic drift, population bottlenecks, eventual introgression between lineages, isolation of populations in fragmented habitats and secondary contacts of previously isolated lineages (Paun et al., 2008; Schneeweiss & Schönswetter, 2011; Voisin et al., 2023), which have led to uneven patterns of inter- and intra-specific diversification in alpine regions today (Schönswetter et al., 2005, 2009; Paun et al., 2008; Alvarez et al., 2009; Schneeweiss & Schönswetter, 2010; Winkler et al., 2010; Aeschimann et al., 2011a; Reisch & Rosbakh, 2021). In particular, during Pleistocene glacial periods, populations with limited ability to disperse often survived locally along the southern Alps by shifting their habitats to different elevations or aspects within the mountainous terrain, resulting in long-lasting genetic isolation that participated in the current high endemism rates observed locally (Tribsch & Schönswetter, 2003; Aeschimann et al., 2011b; Smyčka et al., 2017; Kadereit, 2024). In contrast, populations on the northern side of the Alps, where lowland areas did not allow quick retreat in cold climatic niches at the end of glacial periods, are thought to have experienced a higher extinction rate, and only populations of a few alpine plants, most of them calcicolous since limestone is the most abundant rock type in the northern periphery of the Alps, could survive locally (Andreæ, 1869; Genty, 1932; Thorn, 1960; Tralau, 1963; Kadereit, 2024). This is reflected by a currently high interspecific diversity and intraspecific rarity of genetic lineages in the southern edge of the Alps, whereas the northern Alps nowadays display a high intraspecific diversity, bearing witness to previously extended populations (Gugerli et al., 2008; Aeschimann et al., 2011a; Taberlet et al., 2016; Kadereit, 2024).

The present alpine flora biodiversity is also different in calcareous and siliceous regions. Indeed, siliceous bedrocks occupy the majority of the central Alps, which were mostly covered by ice sheets during extensive glaciations. However, the peripheries are mostly limestone terrains, resulting in more available areas for calcicolous alpine plants to persist during cold periods (Ewald, 2003; Tribsch & Schönswetter, 2003). Therefore, calcicolous species are able to persist locally both during cold and warm periods, resulting in a wider richness for calcicolous species than for silicolous species (Aeschimann et al., 2012). The long-term persistence of calcicolous plants was evidenced in several species. For example, the genetic signal observed in *Androsace lactea* L. showed separated lineages in the southern, the eastern and the northern Alps (Schneeweiss & Schönswetter, 2010). *Erinus alpinus* L. also displayed a diverging genetic pattern in the northern Alps, apart from a migration flux coming from the western Alps to the North (Stehlik et al., 2002). Another species, *Ranunculus alpestris* L. displayed high genetic rarity levels across the southern edge of the Alps, but high genetic diversity across the northern edge, evidencing that large source populations occurred previously in this area (Paun et al., 2008). As a result, the glacial and interglacial history of alpine plants reflects a dynamic process of range shifts, speciation, and extinction (Tralau, 1963; Lang et al., 2023).

The Primulaceae family is composed of over a thousand species, with hundreds adapted to alpine ecosystems. In the Alps, more than 70 species of Primulaceae are present, about a third of which exhibiting strong patterns of endemism, with the highest diversity occurring at subalpine elevations (Aeschimann et al., 2011a,b). This makes the alpine Primulaceae an excellent group for studying the population dynamics of montane plants in relation to climatic fluctuation. For instance, Smyčka et al. (2022) recently demonstrated that time and temperature were the major drivers of diversification in the *Primula* sect. *Auricula*, which mainly occurred in allopatry. The bear’s ear complex (*Primula auricula* L.), consists of a diverse array of taxa, including but not limited to *Primula lutea* Vill., *Primula subpyrenaica* Aymerich, *Primula balbisii* Lehm., and *Primula hungarica* Borbás, many of which have a debated taxonomic status. Due to a possible typification problem, it is currently unclear whether *Primula auricula* L. coincides with *Primula balbisii* Lehm. or *Primula lutea* Vill. In this study, we will refer to *P. auricula* complex in the broad sense as encompassing of all these taxa, mainly growing on the periphery of the Alps and in neighbouring peripheral mountain massifs, and all being specialized to grow on calcicolous substrate – for example limestone, dolomite or conglomerate – (Figure 1). More specifically, the *P. auricula* complex is distributed all across the Alps, except the central part composed of siliceous bedrock (Hartmann & Moosdorf, 2012), as well as in the Pyrenees, the southern Carpathians, the Jura mountains, the Black Forest and the Franconian Jura (Bavaria, Germany) where it forms relictual populations (Schneider & Rudolf, 2013; InfoFlora, 2024). The Jura massif and the unglaciated part – during the Last Glacial Maximum – of the Swiss Molasse Basin (SMB) typically host isolated relicts of mountain plants (Andreæ, 1869; Berthet & Dutartre, 1975; Genty, 1932; Stehlik et al., 2002; Tribsch & Schönswetter, 2003; Kadereit, 2024). These regions are therefore good candidates as potential northern peripheral and lowland glacial refugia (Holderegger & Thiel-Egenter, 2009). In addition, the siliceous central Alps naturally constitute a major geo-physical barrier that could be reflected in genetic differentiation between northern and southern alpine populations.

**Figure 1.**
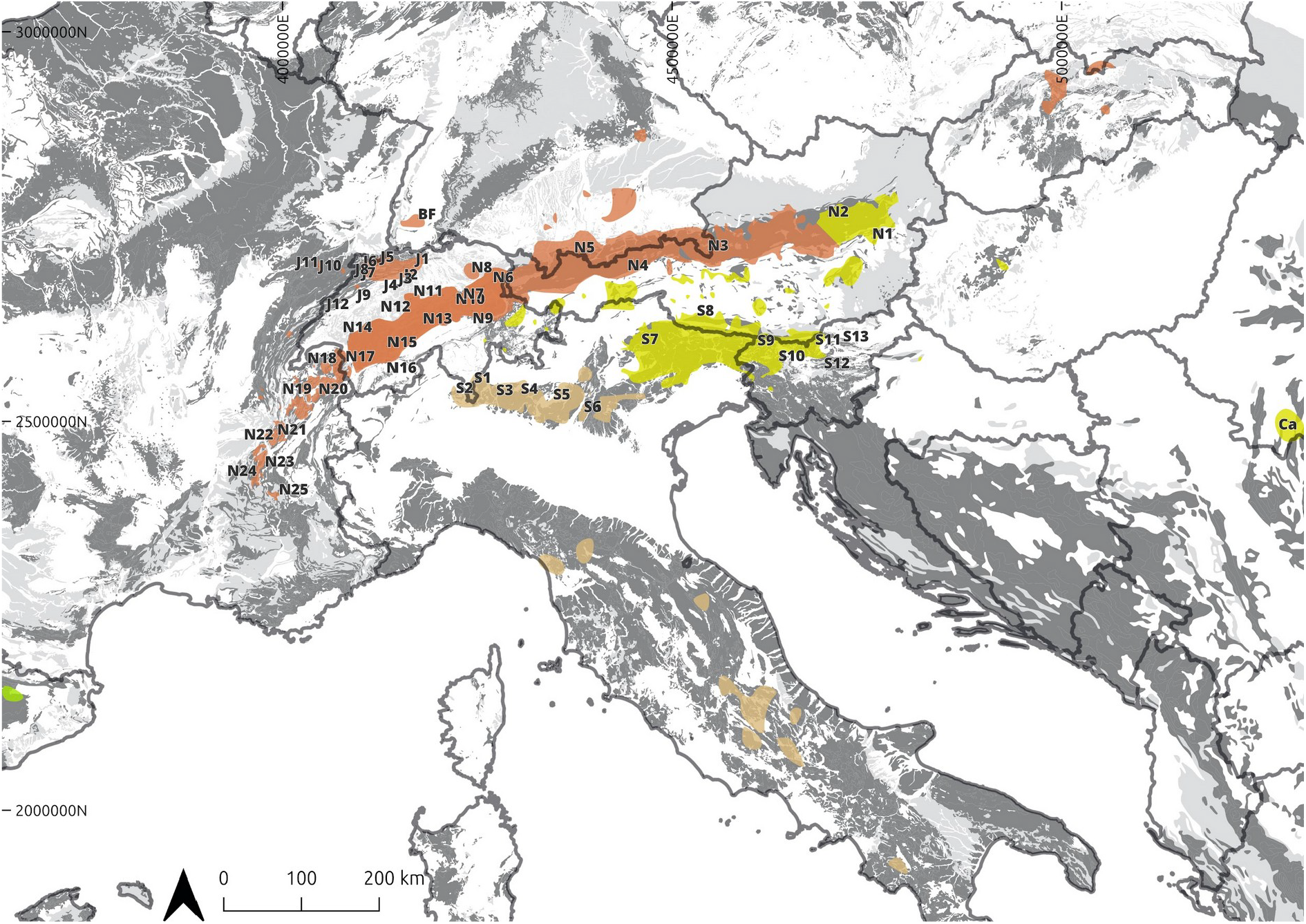
Geographic repartition of the *P. auricula* complex across the European Alpine System. Labels of the sampled localities are mapped across the geographic repartition of *P. auricula*. The northern clade appears in orange (=*P. lutea* Vill.), the south-eastern clade in yellow (=*P. balbisii* Lehm.). *P. subpyrenaica* geographic repartition is displayed in lightgreen and area with unclear taxon identity in camel. The grey shadings represent the calcareous soils according to Hartmann & Moosdorf (2012). Projection : EPSG3035.

Boucher et al. (2016a) summarized the organization of the *P. auricula* complex in the European Alpine System, and highlighted unresolved questions regarding the species’ geographic delimitations in the Lombardian Prealps. Indeed the geographical boundaries between the two lineages present in the Alps could not be precisely defined, particularly because the two taxa that were treated by Boucher et al. (2016a) as separate species (under the name “*P. lutea* Vill.” for the northern clade and “*P. auricula* L.” for the southeastern clade) are morphologically very similar and difficult to distinguish in the field. Past attempts to find discriminating characteristics (L.-B. Zhang & Kadereit, 2004) have sparked literature debate (Somlyay & Bauer, 2010). Here, we used the *P. auricula* complex as a model to explore the potential role of northern peripheral regions of the Alps in diversification processes and the current distribution of calcicolous mountain plants. We specifically investigated: 1) Did peripheral glacial refugia contribute significantly to the genetic diversity observed in the *P. auricula* complex, and how did these populations expand into deglaciated areas after glaciation periods? 2) To what extent have northern peripheral regions such as the Jura massif and the SMB acted as refugia and migration corridors for the *P. auricula* complex? 3) Which were the primary post-glacial recolonization routes followed by lineages within the *P. auricula* complex in the Central European Alps? We hypothesized a) that species in the *P. auricula* complex recolonized the Central European Alps from multiple refugia, including both northern peripheral glacial and southern peripheral glacial refugia, with each lineage following distinct geographic routes shaped by dispersal ability and habitat specialization, b) that populations that recolonized from northern peripheral glacial refugia (e.g., the Jura massif, SMB) exhibit higher levels of genetic diversity compared to populations from southern peripheral refugia, as these peripheral populations were subject to less bottlenecking during glaciation events, and c) that northern peripheral regions, particularly the Jura massif and SMB regions, played a significant role as glacial refugia, enabling calcicolous plants to survive glaciations and later expand into deglaciated areas, thus contributing to the current distribution of these species.

To address these questions, we conducted extensive sampling of the *P. auricula* complex across the Alps and its periphery, including the Jura, the Black Forest and SMB peripheral regions (Tribsch & Schönswetter, 2003; Schönswetter et al., 2005; Aeschimann et al., 2011b; Kadereit, 2024). We performed double-digest Restriction-site Associated DNA sequencing (ddRAD), and applied population genomic approaches and phylogenomic inferences to examine the genetic diversity and delineate the main lineages of *P. auricula* diversity across the Alps, and to assess whether the genetic signal in the northern periphery of the Alps is compatible with a scenario of local survival during glaciations.

## Methods

### Sampling design

Leaf material was collected in 45 localities were sampled across the Alps, Jura, Black Forest and Swiss Molasse Basin. Unfortunately, *P. subpyrenaica* samples could not be included in this study, due to time constraints and unavailability of samples used in previous studies (Boucher et al., 2016a). Each locality was sampled after receiving collection authorizations provided by local governmental offices (Notes S1). Depending on the number of specimens that could be observed in the locality, the topography and the accessibility of the plants, two to five individuals were sampled in each locality respecting a minimum distance of five meters between samples when possible. Fresh leaves were placed in annotated tea bags filled with silicagel and stored at room temperature. Additional material from seven samples of *P. auricula* was received from previous studies (Boucher et al., 2016a) (Table S1).

Five individuals of *Primula spectabili*s Tratt. were included as an outgroup. In total, we sampled 52 localities, 277 individuals of the *P. auricula* complex, and five individuals of *P. spectabilis* across the Alps (Figure S1 and Table S1).

### DNA extraction

Total DNA was extracted from 10-15 mg of dried material, using the Qiagen DNEASY plant mini kit (Qiagen, Hilden, Germany) following the manufacturer’s instructions except for the lysis step which was increased to 50 minutes to diminish secondary compounds’ effect on lysis reaction. DNA concentration was measured using a Qubit 1x HighSensitive assay kit (Thermo Fisher Scientific, Waltham, MS, USA) (see Table S2).

### Library preparation and sequencing

The purified DNA was digested following the ddRAD protocol developed by Peterson et al. (2012), using PstI-HF and MfeI-HF (NEB) enzymes. Digestions were evaluated using a Fragment analyser 5300 (Agilent, Santa Clara, USA). Digested DNA was cleaned using Solid Phase Reversible Immobilization beads (SPRI) (CleanNA) with a ratio of 2:1. P1 and P2 adapters were associated with the digested DNA using T4 DNA ligase. The P1 adapter consisted of 12 different barcodes, the P2 adapter to a common index. The latter was associated with Illumina adapters during the amplification step. Ligated DNA was purified using SPRI beads, using a ratio 1.5:1. The purified ligated DNA was amplified with Platinium SuperFI DNA polymerase (Invitrogen) in a VWR UNO69 Thermocycler (Biometra Göttingen, Germany) in a 32 amplification cycles (72°C during 30 sec) and a final annealing/amplification step (72°C for 5 min). The final product was purified with SPRI beads using a 0.8:1 ratio.

Dual-indexed samples were pooled equimolarly in a final pool. DNA size selection was performed using a Pippin Prep 2% DF agarose cassette for size selection ranging between 250-300 bp (Sagescience), and concentrated thanks to Microcon 10K centrifugal filter device (Merck). In total, two libraries were sequenced on single-end Illumina Nextseq 500 Instrument, with High-Output Flow cell and 15% PhiX at Fasteris sequencing facility (Geneva).

### Sequence filtering and alignment

Demultiplexing was performed in Cutadapt 4.4 (Martin, 2011). We allowed no indels and N across the sequences. Data quality trimming on the 3’ tail was performed on Ipyrad v.0.9.92 (Eaton & Overcast, 2020), allowing a maximum of 5 low quality bases with (max_low_qual_bases = 5), and a minimum quality-score (Qscore) set to 33 (Phred_Qscore_offset = 33). Reads were clustered into shared locus using Ipyrad (Eaton & Overcast, 2020), and bad quality samples were filtered following Cerca et al. (2021). After removing 48 bad-quality samples, a set of 229 samples and five outgroup samples of *P. spectabilis* were obtained. The best clustering threshold was assessed by comparing statistics of cumulative variance explained by the eight first axes of PCA and by the correlation between genetic distance and missingness of seven threshold values from 0.84 to 0.96 stepping 0.02 (McCartney-Melstad et al., 2019). The best clustering threshold, i.e. 0.90 ensured a high proportion of explained variance in the PCA associated with a low correlation between genetic distance and missingness. The resulting loci were used for phylogenetic inference and datation, while SNPs were used for population genetic analysis.

### Phylogenetic inference and dating

Loci were aligned using the Mafft v.7.505 program with auto option (Katoh & Standley, 2013), resulting in a total of 25,660 loci. The loci filtering threshold was assessed by comparing the phylogenetic inferences statistics. Particularly, the weighted mean bootstrap support was evaluated thanks to a balanced index calculation, detailed in (McCartney-Melstad et al., 2019), which consists in multiplying bootstrap support of each node in the consensus trees by the number of their child nodes.

By doing so, the loci filtering threshold was fixed to 50 % of the total samples resulting in a library of 1688 loci. The gap between the initial number of loci retrieved and the filtered loci was explained by the fact that many retrieved loci in the initial library consisted in private loci to unique samples, to loci retrieved only in *P. spectabilis*, or only in a few samples. The library was concatenated in phylip files using the SEGUL v.0.22.1 and Goalign v.0.3.7 program (Handika & Esselstyn, 2024; https://github.com/evolbioinfo/goalign). The best partition schemes were determined using PartitionFinder.2.1.1 (Lanfear et al., 2017) and phylogenetic inferences were identified using IQ-TREE 2 2.2.2.6 (Minh et al., 2020), with best fit model testing followed by direct tree construction (option -m MFP), using UFBoot trees optimization (-bnni option), 1000 replication tests for assessing bootstrap support (Hoang et al., 2018) and branching support (SH-aLRT test) (Guindon et al., 2010).

The divergence between *P. spectabilis* and the *P. auricula* complex has been estimated to 6.6013 Mya (Boucher et al., 2016b). Therefore, the root of the dating inference, consisting in the divergence between *P. spectabilis* and the *P. auricula* complex was forced accordingly. The dating was performed using IQTREE 2 2.4 and 10000 replicates. Since they rely on secondary calibration, cautious interpretation of these results can be used to give an idea of divergence dynamics inside the *P. auricula* complex in the Alps.

As a complementary approach to the concatenation-based maximum likelihood tree inferences, a coalescent species-tree approach was performed. Loci containing no informative sites for parsimony were filtered using SEGUL (Handika & Esselstyn, 2024) with align filter subcommand (option --pinf 1). Individual gene trees were inferred for each of the remaining 1,611 loci using IQTREE2 2.2.2.6 (Minh et al., 2020) with best fit model testing followed by direct tree construction (option -m MFP), using AICc optimality criterion, and 1000 SH-aLRT replicates. In order to reduce gene tree estimation error that can arise when resolving hundreds or more clades using short coalescent-gene alignments with few parsimony-informative characters, clades with 0% SH-like aLRT support were collapsed following Simmons & Gatesy (2021). The species tree was estimated by combining gene trees using ASTRAL-IV v1.20.4.6 (C. Zhang & Mirarab, 2022) by optimizing the objective function of ASTRAL (C. Zhang et al., 2018). Branch lengths were computed using integrated CASTLES-II (Tabatabaee et al., 2023).

Using the ML phylogenetic tree as guide tree and the gene trees rooted on outgroup samples using Phyx (Brown et al., 2017), the trinomial distribution of triplets (tr2) (Fujisawa et al., 2016) method was used to investigate species delimitation and distinct evolutionary lineages among the genetic clusters.

### Ancestry range estimation (ARE)

The ancestral range on each node of the ML tree after removing outgroups was estimated using a Bayesian approach implemented in the R package ‘phytools’ (Revell, 2024). Eight geographical areas where the *P. auricula complex* occur were defined according to main geographical barriers across the Alps (Figure 2a): (A) Lombardian Prealps and the Ticino region to the Monte Baldo (Italy) and (B) the southeastern Alps were separated by the Adige valley; (C) the southern Carpathians and (D) the northeastern Alps to the Rhine valley were separated by the siliceous regions in central Alps and pannonian basin; (E) the northern prealps and SMB were separated from (F) the western and southwestern Alps by the Rhône valley; (G) the Black Forest was separated from the (H) Jura massif by the Rhine valley. We did not allow one lineage to occupy two regions simultaneously. Three unordered evolutionary models were built using the *fitMk* function : the Equal Rate (ER), the All Rate Different (ARD) and the Symmetric Model. The models were compared with an ANOVA, based on the Akaike Information Criterion (AIC). The ER model, which displayed the lower AIC index, was selected for further 1000 stochastic simulation mappings (function *make*.*simmap*) and final ancestral range estimation on nodes.

**Figure 2.**
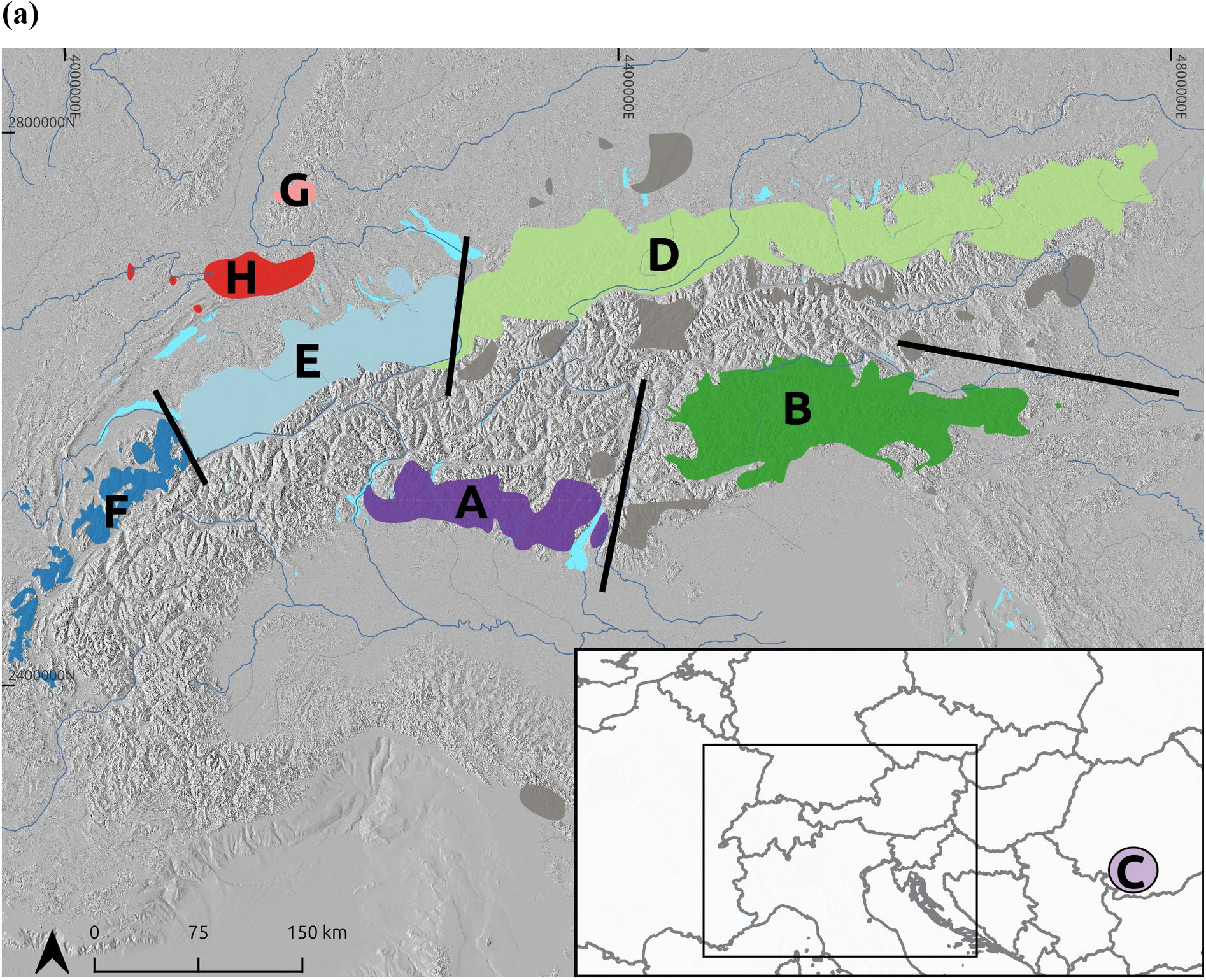
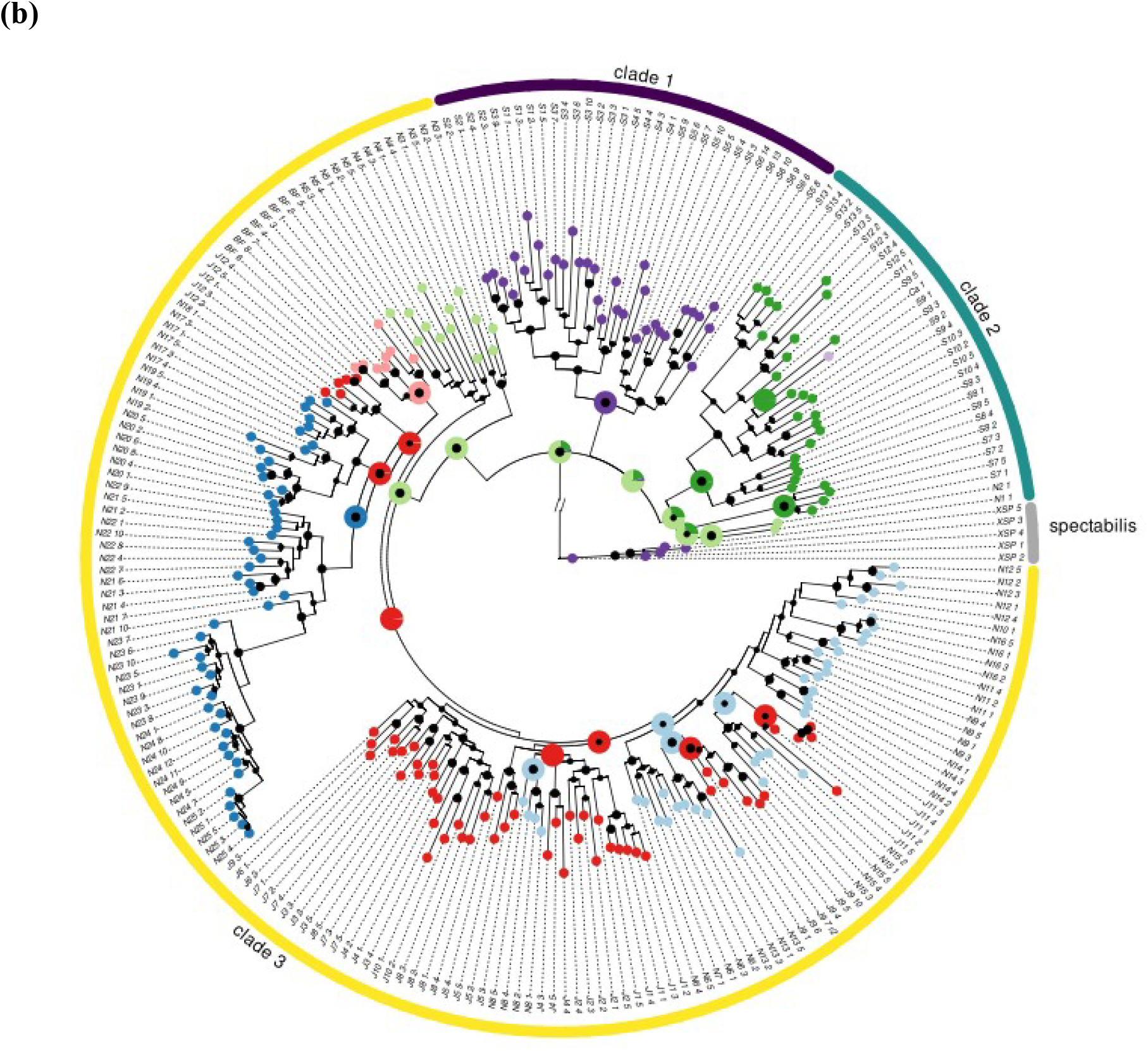
(a) Definition of the geographic regions where the *P. auricula* complex occurs across the Alps. (b) Maximum-Likelihood phylogeny with ancestry range estimation on nodes of the *P. auricula* complex across the Alps. Violet, blue and yellow colours represent Clades 1, 2 and 3 respectively, while grey branches correspond to *P. spectabilis* samples. Big black circles indicate node support with SH-aLRT >80 and UF-Boot >90, smaller black circles circles indicate node support with SH-aLRT >80 or UF-Boot >90. Pie charts on nodes represent the ancestral range estimations (ASE).

### Population structure

The raw variant calling file from Ipyrad was filtered with VCFtools v.0.1.16 (Danecek et al., 2011) to keep bi-allelic SNPs shared by at least 50% of the samples and to remove indels. For the population cluster analysis, Minor Allele Frequency threshold (--maf) was set to 0.05 and one SNP by locus was randomly extracted to keep unlinked SNPs. Outgroup samples were removed to focus on the *P. auricula* complex population structure. Snmf was used to assess population admixture across the whole distribution of the *P. auricula* complex (Porras-Hurtado et al., 2013; Frichot et al., 2014). The Cross-Entropy criterion was calculated and compared across ten replicates for each K value from 1 to 10. Clustering was assessed based on the Cross-Entropy index.

### Population-level statistical analyses

Population statistics were estimated using the ‘hierfstat’ R package (Goudet et al., 2015). Observed heterozygosity (Ho) was estimated for the main genetic clusters identified in the phylogeny and the population structure results. Pairwise differentiation indices (FST) were estimated with the *pairwise*.*WCfst()* function from the ‘hierfstat’ R package (Goudet et al., 2015) across the main genetic groups. The individual inbreeding coefficient (FIS) was calculated in each sampled locality with the --het parameter in VCFtools v.0.1.16 (Danecek et al., 2011). Finally, to assess potential isolation by distance (IBD), Mantel tests were performed at the sampled locality level between genetic distance matrices obtained previously and geographic distance matrices using the ade4 R package (Dray & Siberchicot, 2017).

## Results

DNA concentration before the PCR step reached an average of 6.33 ng/µl (sd = 5.24), the concentration after the PCR step reached an average of 8.32 ng/µl (sd = 5.91), and Illumina sequencing yielded a total of 495,929,307 reads with a mean of 2,101,395 reads per sample (detailed in Table S2). The final dataset consisted of 229 individuals of the *P. auricula* complex and five of *P. spectabilis* containing less than 30 % missing data, 1.688 loci shared across at least 50 % of the total samples. These loci comprised 225,354 sites, of which 127,511 were variant after filters, or 3,489 variant sites when the Minor allele Frequency was set to 0.05.

### Phylogenetic inference

The Maximum likelihood phylogeny (ML tree) was rooted manually to separate the specimens of *P. spectabilis* and the *P. auricula* complex (SH-aLRT = 100 and UF-Boot = 100). Samples from the same locality generally formed a clade and the mean bootstrap support reached the value of 83.76% with 8201 parsimony-informative sites, 3,330 singleton sites and 213,823 constant sites according to the log file from IQ-TREE command.

The rooted tree displayed homogenous geographic organization, as samples from a same locality, and localities from the same massif formed monophyletic groups (except a few exceptions in the northern Alps). The first supported split inside the *P. auricula* complex differentiated three main clades inside the *P*.*auricula* complex. Clade 1 included samples from geographic region A (SH-aLRT = 100 and UF-Boot = 100) (Figure 2). Inside this clade, the Lugano and Bergamasque Prealps (SH-aLRT = 100 and UF-Boot = 100), and the Brescia and Garda Prealps separated in two subclades (SH-aLRT = 99.9 and UF-Boot = 100). Clade 1 and Clade 2 separated near the Adige Valley. Clade 2 extended further east and north and reached regions B, C and the northeastern limit of region D (SH-aLRT = 98.3 and UF-Boot = 90). Inside Clade 2, the Dolomitic Alps, the Styrian Prealps and the northern Lower Austria Alps are initially separated from the rest of the samples (SH-aLRT = 93.8 and UF-Boot = 89). Among the remaining samples, the western Tauern Alps formed another isolated cluster (SH-aLRT = 100 and UF-Boot = 100) followed by two subclades. One included the Julian Prealps and the carpathian locality (SH-aLRT = 99.4 and UF-Boot = 99), the second included the Slovenian and Carinthian Prealps and Alps (SH-aLRT = 96.5 and UF-Boot = 93) (Figure 2). Clade 3 extended from regions D, E, F, G and H (SH-aLRT = 97.5 and UF-Boot = 100) (Figure 2). In Clade 3, strong support (SH-aLRT = 100 and 92.9 and UF-Boot = 100 and 78) was found for the divergence between the easternmost populations, situated in the Berchtesgaden, the Bavarian and the northern Tyrolian Alps and the remaining localities which diversified in two groups. Region G was clustered with region F and the southernmost geographical limit of region H, forming an “external” clade (Figure 2) (SH-aLRT = 99.1 and UF-Boot = 91). Particularly in this group, region F was connected by a branch much longer than any other within Clade 3. The remaining localities, situated in regions E and H formed an “internal” clade (SH-aLRT = 60.3 and UF-Boot = 64).

The tree computed with Astral IV from individual gene trees displayed a different organization on the deepest branches. Clade 2 described in the ML tree topology was the first group to diverge inside the *P. auricula* complex (Figure S2). However, the Styrian Prealps and the northern Lower Austria Alps did not belong to this clade and formed a sister clade of Clades 1 and 3. Inside Clade 3, populations from the Berchtesgaden Alps remained the earliest diverging lineage. However, the “external” and the “internal” clades did not display the same organization as in the ML tree. Region F was the second clade to split up. Region G and the southernmost locality from region H set up the basis of further diversification in the rest of region H as well as in region E.

In the next sections, Clade 1, Clade 2 and Clade 3 will refer to the clades described in the ML tree.

### Ancestry range estimation (ARE)

The Equal Rate (ER) model of evolution best supported the dataset (AIC = 157.28, weight = 1) compared to the ARD (AIC = 260.52, weight = 3.815e-23) and to the SYM (AIC = 205.38, weight = 3.595e-11) models of evolution. Results from this analysis suggested that the *P. auricula* complex ancestor originated near the Clade 2 geographical range, and in particular from the northeastern geographical limit of region D. According to the selected model, Clade 3 also originated from the same region, the basal nodes splitting the populations in an east-west direction from the Bavarian and the northern Tyrolian Alps to the Berchtesgaden Alps. Deeper in the branches of Clade 3, the analysis recovered the Jura mountains (region H) as the ancestral area/region shared by all populations from the Jura mountains (region H), Black Forest (region G) northern Alps *sensu stricto* (region E) and western Alps (region F).

### Species delimitation and population structure

The species delimitation analysis (using tr2) separated three distinct evolutionary lineages across the Alps, corresponding to clades 1, 2 and 3 as described in the ML tree. (Figure 3). Moreover, the best clustering result with the sNMF software was obtained for K=4 (Figures 3-4). Two clusters matched the geographic delimitations of Clades 1 and 2. In Clade 3, two clusters separated region F from the rest of the samples (Figure 2a, Figure 4). Localities from the western limit of region B and northeastern limit of region D as well as the northernmost localities of region F (Figure 2a) displayed a mixed genetic ancestry. When increasing the number of ancestral populations, the clustering pattern aligned progressively on the clades retrieved by the ML phylogenetic topology (see Figure S3). Finally, the population admixture analysis of the *P. auricula* complex across the Alps demonstrated a clear geographical structure. The Mantel correlation test showed a significant correlation between geographic distance and genetic distance between samples (r = 0.559, p-value < 0.001).

**Figure 3.**
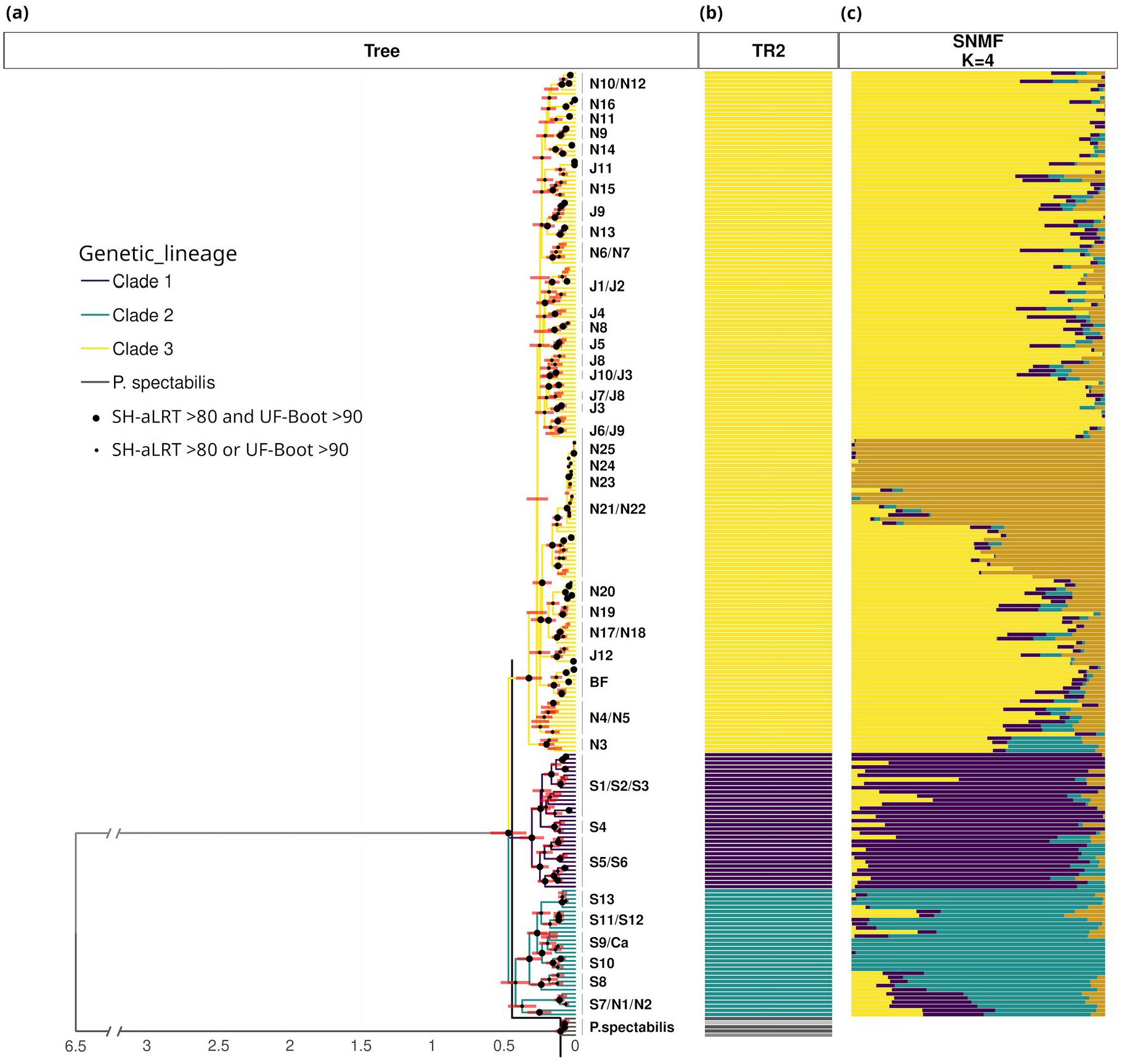
Results of divergence estimation, species delimitation and admixture proportions in the *Primula auricula* complex across the Alps. **(a)** The ML tree is scaled according to the estimated divergence times. The time scale (in Mya) is added as x axis, and the dating intervals for each estimated node are added as red bars on each node. Violet, blue and yellow colours represent Clades 1, 2 and 3 respectively, while grey and lightgreen branches correspond to *P. spectabilis*. Big black circles indicate node support with SH-aLRT >80 and UF-Boot >90, smaller black circles indicate node support with SH-aLRT >80 or UF-Boot >90. **(b)** The result of the species delimitation (TR2) is represented with colored bars. The grey, green, violet, blue and yellow separate the distinct lineages defined by the software. The black bar on the tree illustrates the delimitation. **(c)** Cumulative barplots representing the population admixture computations evidenced with the sNMF software. Admixture proportions are colored according to the deepest curve of cross-entropy index among k clusters, K=4.

**Figure 4.**
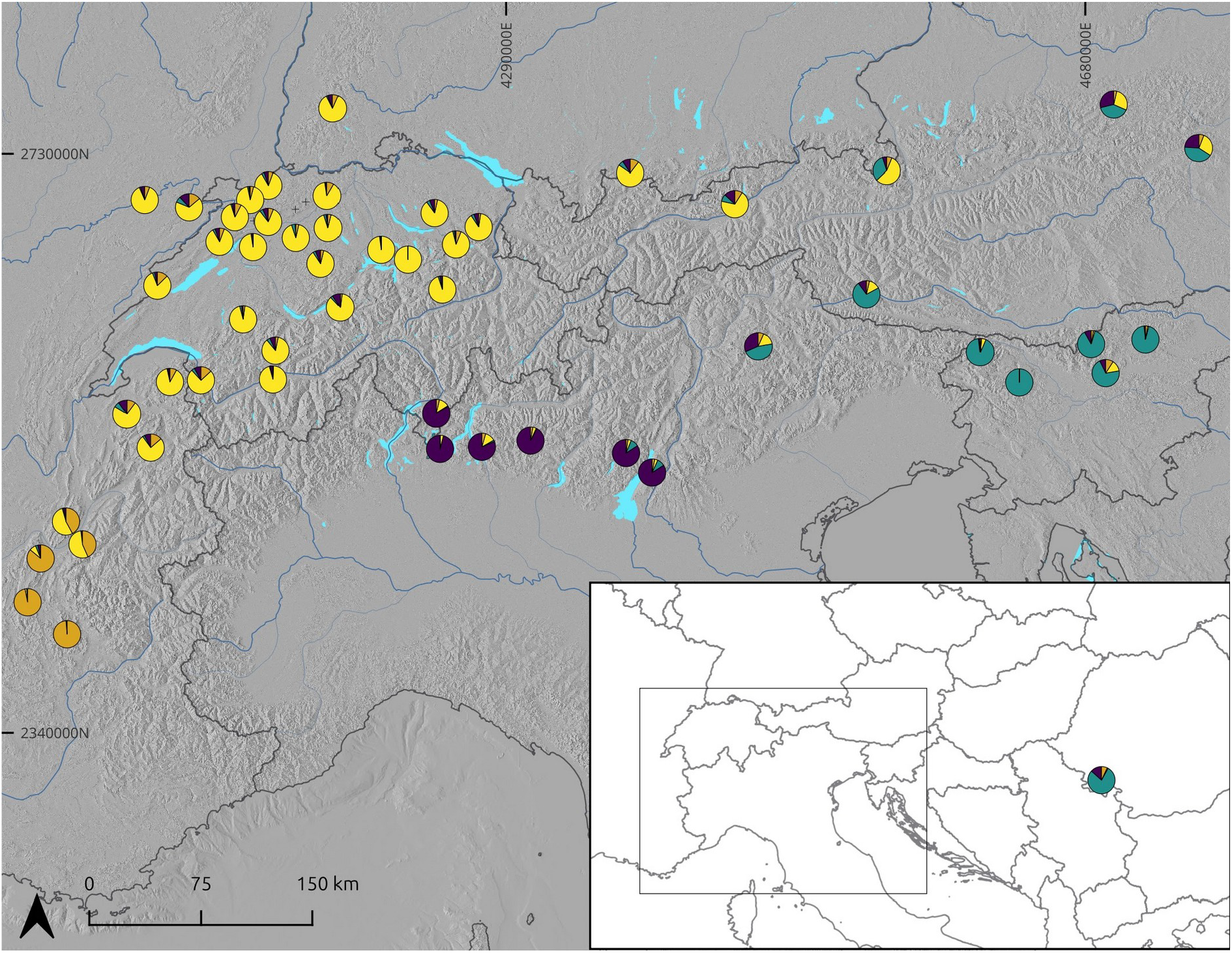
Admixture pattern of the *P. auricula* complex across the Alps from sNMF analysis for K=4. The colored diagrams correspond to the parental lineage admixture identified by the software for each sampled locality. Map background : GEBCO Bathymetric Compilation Group 2022 (2022) Projection: EPSG3857

### Divergence time estimation

The three clades described in the ML tree diverged near 0.47 Mya (± 0.14 - 0.10) (Figure 3). Diversification inside Clade 2 separating the Dolomitic, the northern Lower Austrian and the Styrian Alp localities from the eastern region of the Piave valley including the Carpathian locality, occurred later, approximately 0.37 Mya (±0.13-0.07). The western Tauern Alps as well as the Berchtesgaden Alps from Clade 3 and the divergence between the splitting the Lugano and Bergamasque Prealps from the Brescia and Garda Prealps form Clade 1 occurred in similar times (0.30 Mya, error : ~ 0.05 Mya). Further recent diversification events inside each Clade occurred during the Mid and Late Pleistocene. In particular, populations from the French prealps diversified recently, between 0.23 (± 0.09-0.05) and 0.16 Mya (±0.05,0.03).

### Population level descriptive statistics

Clade 3 described in the ML tree exhibited lower observed heterozygosity (Ho = 0.0642) in comparison to Clade 2 (Ho = 0.0815) and Clade 1 (Ho = 0.072). The mean allelic richness in Clade 3 was lower (mean = 1.136, 3rd quantile = 1.233), compared to Clade 1 (mean = 1.144, 3rd quantile = 1.264) and Clade 2 (mean = 1.170, 3rd quantile = 1.333). The mean value of the inbreeding coefficient (Fis) across loci in the total sampling area (0.588) and in each Clade (between 0.45 and 0.63) was very high indicating an overall high inbreeding proportion in groups. The pairwise-Fst value between Clade 1 and Clade 2 (0.272) was slightly lower than the pairwise-Fst values between Clade 3 and Clade 1 or 2 (0.303 and 0.297 respectively).

## Discussion

The present study offers a global view on the geographic structure of the *P. auricula* complex across the Alps. By combining phylogenetic approaches and population genomics, we reconstructed the evolutionary history of lineages and assessed the current structure of populations.

### Phylogeographic structure of the *P. auricula* complex across the Alps

According to the dated phylogenetic inferences and to the ancestry range analysis, the *P. auricula* complex likely diversified in three well defined clades during the late Miocene (Figure 3). Two of them, Clade 1 and Clade 2, are separated from west to east, near the Adige valley running from Innsbruck (AU) to the Garda lake (IT), a well-recognized geographical barrier at the community level (Chodat & Pampanini, 1902; Thiel-Egenter et al., 2009). Clade 1 comprises essentially populations of the southern Alps, from both the Lombardian Prealps and Ticino regions, recognized as the highest center of endemism in the Alps (Aeschimann et al., 2011a; Kadereit, 2024; Parisod, 2022). This region was not included in previous studies about the *P. auricula* complex, except for one genetically isolated individual from the Monte Baldo locality, interpreted as a hybrid zone between Clade 2 and Clade 3 (described as *P. auricula* and *P. lutea* in Boucher et al. (2016a)). Instead of a hybrid zone, by further sampling in this area, the present study unravels the existence of a third clade in the *P. auricula* complex across the Alps *sensu lato*. Clade 2 groups all populations from the eastern Alps from Trentino to the northeastern Alps and includes the population from the southern Carpathians. Clade 3 covers all the northern Alps from the Berchtesgaden Alps to the western Alps, as well as the surrounding peripheral areas: the Jura massif, the Black Forest and the Swiss Molasse Basin (SMB). This group is geographically extensive and genetically homogenous compared to Clade 1 and Clade 2. Meanwhile, the extensive sampling in this area highlights recent diversification dynamics, including strong genetic divergence at the periphery of the distribution of this complex. Regional isolation from a hypothetic northern lowland and peripheral glacial refugia were also revealed thanks to ARE between the geographic regions included in this area. (see Figure 2).

### A Geographical fragmentation in the southern peripheries of the Alps

The southern Alps are made of jagged mountains, bounded by deep U-shaped valleys that act as strong physical barriers, delineating never glaciated areas (Ehlers & Gibbard, 2011), and making this region the highest center of endemism across the Alps (Aeschimann et al., 2011a; Parisod, 2022; Kadereit, 2024). This pattern of multiple fragmentation is recovered in the *P. auricula* complex which displays strong splitting scheme in both Clade 1 and Clade 2, separating the southern Alps sampled localities in geographically restricted subclades supporting a long term geographical isolation *in situ*. These subclades overlapp putative peripheral glacial refugia proposed by Tribsch & Schönswetter (2003) and the time-divergence analysis performed in the present study dated this split between 0.62 and 0.37 Mya, corresponding to the early mid-quaternary climatic fluctuations (Ehlers & Gibbard, 2011). The pattern observed in this study on the southern Alps is in accordance with previous literature focusing on other low-mobile organisms (Scheel & Hausdorf, 2012), and adds supplementary support on both glacial and interglacial isolation in southern peripheral refugia of the Alps (Kadereit, 2024).

### Gene flow across the eastern Alps challenges physical barriers

The southeastern Alps region is particularly interesting for calcicolous plant phylogeography. Genetic flow between disjunct calcareous areas of the southeastern and the northeastern Alps was demonstrated in calcicolous mountain plants, for example *Androsace lactea* L. (G. Schneeweiss & Schönswetter, 2010), *Papaver alpinum* L. (Schönswetter et al., 2009) or *Ranunculus alpestris* L. (Paun et al., 2008).

Similar genetic signal is retrieved in this study, as the Styrian Prealps and the northern Lower Austrian Alps and the Carpathian locality from Clade 2 show low genetic differentiation between the southeastern and northeastern Alps. The carpathian locality, the southeastern and the northeastern Alps are separated by intermediate siliceous regions, a strong ecological barrier for strictly calcicolous plants such as the *P. auricula* complex. While a putative glacial refugia for calcicolous mountain plants is supported by recent literature in the deglaciated eastern Pannonian basin composed of sedimentary rocks (Slovák et al., 2012), the persistence of mountain calcicolous plants in this region cannot explain the ability of such species to cross the strong siliceous barrier between regions, unless considering long-distance dispersal.

### Danubian pathway hypothesis

In the northeastern Alps, historical observations relate the presence of the *P. auricula* complex near Danubian water beds (Essl, 1993) as well as in peatland areas of the Munich gravel plain (Bräuchler et al., 2015). It can still be observed today along the Danube valley in the Kelheim locality, Bavaria, Germany (SM, personal observation). Short distance or long distance dispersal in such habitats through abiotic or biotic factors (water, wind, ants for example) could allow plant species to cross physical barriers and permit *P. auricula* complex exchanges between the Alpine arc *sensu stricto* and the peripheral massifs (Casazza et al., 2008, Anjos et al., 2020). A closer look at this region is particularly interesting to deepen the diversification history of Clade 3. The Franconian Jura and northern lowland areas remained unglaciated during the Pleistocene and were hypothesized as lowland glacial refugia for several species (Tremetsberger et al., 2002). According to the results given by the ARE performed in this study (Figure 2), this region might have driven an allopatric diversification in the *P. auricula* complex followed by migration after glaciers retreat.

### Different latitudes, different dynamics

The presence of the *P. auricula* complex across all of the Alps and peripheries makes it a good model to compare phylogeographic dynamics between the southern and northern parts of the Alps. Populations in the southern Alps (Clades 1 and 2) remained geographically separated and show increased genetic differentiation, leading to a low allelic richness (1.144 and 1.170 respectively,) associated with high heterozygosity. The contrasting northern Alps lineage (Clade 3) displays genetic homogeneity with a lower observed heterozygosity (0.064) and allelic richness (1.136) compared to Clade 1 and 2 associated with a deficit of heterozygotes (mean Fis = 0.588). This homogeneous genetic group, compared to the fine genetic structure observed in Clades 1 and 2 from the southern and southeastern Alps evidences contrasting historical dynamics, likely including extended hypothetical ancestral populations and recent gene flow, supported by the ARE (Loveless & Hamrick, 1984; Ibrahim et al., 1996; Parisod, 2022; Kadereit, 2024). These results are in line with the general literature on alpine plants, which generally classify this region as an area with low-to-null species and genetic endemism (Aeschimann et al., 2011a), but a high allelic richness compared to the southern Alps (Taberlet et al., 2016). As suggested in Kadereit (2024), allopatry in this region may only happen during interglacial periods which were mainly shorter in time compared to glacial periods in the past, and did not allow for the completion of the speciation process. This dynamic process of isolation and recolonization in the *P. auricula* complex might also explain the contrasting results observed on allelic richness compared to previous literature. However, further efforts should deepen the understanding of fine-scale dynamics in this area.

### Limitations of the phylogenetic approach and perspectives

The results obtained in this study confront the limitations of the phylogenetic and clustering approach to understand complex phylogeographic stories of mountain plants. Particularly, these methods could not allow to disentangle which dynamics shaped the local identities in intermediate regions between big geographic regions defined in the ARE analysis. Both the Dolomitic Alps, the Styrian Prealps, the northern Lower Austria and the Bavarian Prealps could be remnants of old lineages, as suggested by their basal positions in Clade 2 and Clade 3 (Figure 2b). However, each of these regions are situated at the external geographic limit of region B (Figure 4). Consequently, these regions also constitute potential hybrid zones between Clade 1 and Clade 2, and Clade 2 and Clade 3, an idea which is supported by our clustering approach with sNMF. A denser sampling in intermediate regions should offer new clues to disentangle evolutionary processes in such areas.

Moreover the young and rapid phylogeographic dynamics in Clade 3 might create incongruence between topologies in ML tree and gene tree, creating incomplete lineage sorting effects and possible past hybridization imprints between fluctuating groups. For example the Dauphiné prealps included in Clade 3 evidence strong genetic differentiation. However, the approaches used in the current study do not allow to disentangle this pattern between processes such as a geographic isolation with a strong bottleneck, migration of few individuals or putative hybridation with a ghost lineage, for example the pyrenean *P. subpyrenaica* Aymerich, Sáez & López-Alvarado (Slatkin & Takahata, 1985), each of these processes leading to long branches in phylogenies (Hibbins & Hahn, 2022). Particularly, extensive samplings in other peripheral massifs of the Alps such as the Apennines, the Dolomites or the Tatras Mountains might deepen the dynamics evidenced in this research, as well as including the pyrenean population of the recently described endemic *P. subpyrenaica* (Aymerich et al., 2014), a sister-species of Clade 3 from the *P. auricula* complex included in this study according to (Boucher et al., 2016a). Overall, phylogeographic studies on alpine plants should broaden their area of interest when considering sampling designs, as peripheral massifs relate an entire chapter of their life: the glacial history of such species.

## Data accessibility

Raw demultiplexed genetic sequences are available at: Accession number: PRJNA1245062; https://www.ncbi.nlm.nih.gov/sra/PRJNA1245062 (detailed in Table S3)

The general bioinformatic pipeline of the study is available on the Public GitLab repository primula_phylogenetics v1.0.0 : https://gitlab.com/phylo/primula_phylogenetics.git

