## Supplementary_information for "Peripheries in alpine phylogeography"

**Supporting Information**

**The role of peripheral regions in shaping the phylogeography of the Bear’s ear complex across the Alps**

**Figure S1** Specimens from the *P. auricula* complex in their natural environment, and sampling localities across the Alps.

**Figure S2** Species tree reconstruction of the *P. auricula* complex across the Alps, with weighted-ASTRAL

**Figure S3** Population admixture for the *P. auricula* complex across the Alps, based on the sNMF software computations. Admixture are colored among k clusters, top : K=5, middle : K=6, bottom : K=7.

**Table S1** Sampled localities and collectors of the P*. auricula* complex across the Alps

**Table S2** DNA concentrations of *P. auricula* samples before PCR amplification and ipyrad library statistics

**Table S3** *Primula auricula* and *Primula spectabilis* demultiplexed sequences: NCBI availability

**Note S1**  Collection authorizations for the *Primula auricula* samples in the Alps.

**Figure S1** Specimens from the *P. auricula* complex in their natural environment: (a) at Le Quénet locality (J6), Switzerland (Philippe Juillerat) ; (b) at Chasseral locality (J9), Switzerland (Stéphanie Morelon) ; and sampling localities across the : (c) European geographic distribution of *Primula auricula* complex. The sampled areas are indicated by black boxes. The Carpath sampling consisted of one locality (labelled Ca). The locality is indicated by the black box ꞵ. The Alps sampling of the box α. is detailed in (d), indicating the locality labels.

The current occurrences of Primula auricula complex were obtained from Infoflora (CH), Conservatoire Botanique de France (Alpin and Franche-Comté), Glick Fridjof (GE), and a strict selection of occurrences from GBIF dataset.

**Figure S2.** Species trees of the P. auricula complex and outgroups (P. spectabilis) built by combining locus trees using the weighted-ASTRAL optimization algorithm, and taking into account phylogenetic uncertainty by relying on branch length and branch support across locus trees.

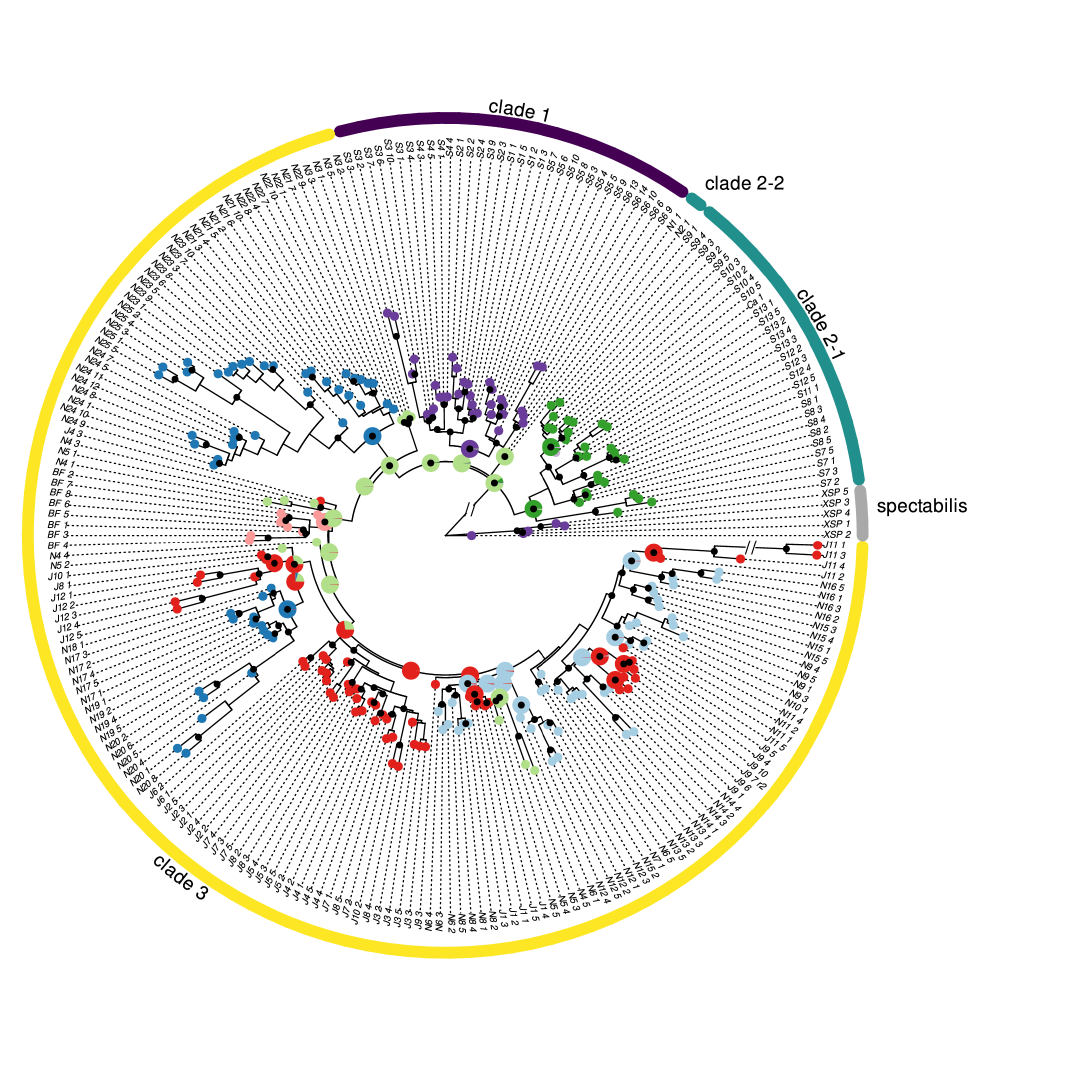

**Figure S3** Population admixture for the P. auricula complex across the Alps, based on the sNMF software computations. Admixture are colored among k clusters, top : K=5, middle : K=6, bottom : K=7.

**Table S1** Sampled localities and collectors of the *P. auricula* complex across the Alps

| **Locality Name** | **Locality Label** | **X4326** | **Y4326** | **Coordinates precision (meters)** | **Collector.s** | **Number of  collected individuals** |
| --- | --- | --- | --- | --- | --- | --- |
| CH_Denti_Vecchia | S1 | 46.05352 | 9.02327 |  | Philippe Juillerat | 5 |
| CH_Generoso | S2 | 9.02177 | 45.92935 | 16 | Mariama Diallo, Thomas Bulliard et Stéphanie Morelon | 5 |
| IT_Grigna | S3 | 45.92169 | 9.38959 |  | Thomas Bulliard et Philippe Juillerat | 10 |
| IT_Pizzo_Arera | S4 | 45.93113 | 9.80327 |  | Thomas Bulliard | 5 |
| IT_Monte_Stigolo | S5 | 45.85149 | 10.63445 |  | Thomas Bulliard et Philippe Juillerat | 10 |
| IT_Monte_Baldo | S6 | 45.72395 | 2388231 |  | Adrian Möhl | 10 |
| IT_Punta Penia | S7 | 46.49247 | 11.81033 |  | Thomas Bulliard | 5 |
| AT_Kreidenfeuer | S8 | 46.80239 | 12.75676 |  | Thomas Bulliard | 5 |
| SI_Kraj_sten | S9 | 46.42665 | 13.75241 |  | Thomas Bulliard | 5 |
| SI_Altemaver | S10 | 46.23376 | 14.09137 |  | Thomas Bulliard | 5 |
| SI_Robanov_Kot | S11 | 46.3983 | 14.7072 |  | Thomas Bulliard | 5 |
| SI_Veliki_vrh | S12 | 46.25587 | 14.84185 |  | Thomas Bulliard | 5 |
| SI_Tisnik | S13 | 46.41319 | 15.17492 |  | Thomas Bulliard | 5 |
| RO_Baile Herculane | Ca | 44.8838 | 22.42909 |  | Florian Boucher-Gabriele Casazza-Peter Szovenyi-Elena Conti - Molecular Phylogenetics and Evolution 104 (2016) 60:72 | 1 |
| AT_Raxalpe | N1 | 47.575 | 15.79167 |  | Florian Boucher-Gabriele Casazza-Peter Szovenyi-Elena Conti - Molecular Phylogenetics and Evolution 104 (2016) 60:72 | 1 |
| AT_Lunz_am_See | N2 | 47.8685 | 15.05913 |  | Florian Boucher-Gabriele Casazza-Peter Szovenyi-Elena Conti - Molecular Phylogenetics and Evolution 104 (2016) 60:72 | 1 |
| DE_Berchtesgaden | N3 | 12.94583 | 47.57593 | 0 | Philippe Juillerat et Stéphanie Morelon | 5 |
| AT_Innsbruck | N4 | 11.6 | 47.3 |  | Pau Carnicero | 1 |
| DE_Füssen | N5 | 10.75759 | 47.56056 | 7 | Philippe Juillerat et Stéphanie Morelon | 5 |
| CH_Säntis | N6 | 9.31946 | 47.22256 | 2.5 | Philippe Juillerat et Stéphanie Morelon | 5 |
| CH_Amden_Gulmen | N7 | 47.1679 | -42853 |  | Florian Boucher-Gabriele Casazza-Peter Szovenyi-Elena Conti - Molecular Phylogenetics and Evolution 104 (2016) 60:72 | 1 |
| CH_Tössbergland | N8 | 8.95244 | 47.33446 | 3 | Philippe Juillerat et Stéphanie Morelon | 5 |
| CH_Linthal | N9 | 8.97979 | 46.88237 | 2.7 | Philippe Juillerat et Stéphanie Morelon | 5 |
| CH_Mythen | N10 | 47.02972 | 1822410 |  | Florian Boucher-Gabriele Casazza-Peter Szovenyi-Elena Conti - Molecular Phylogenetics and Evolution 104 (2016) 60:72 | 1 |
| CH_Rigi | N11 | 8.97979 | 46.8823 | 2.7 | Philippe Juillerat et Stéphanie Morelon | 5 |
| CH_Napf | N12 | 7.94803 | 47.00705 | 8 | Philippe Juillerat et Stéphanie Morelon | 5 |
| CH_Isetwald | N13 | 8.14575 | 46.72941 | 3.5 | Philippe Juillerat et Stéphanie Morelon | 5 |
| CH_Baderhore | N14 | 7.32059 | 46.612499 |  | Thomas Bulliard | 5 |
| CH_Wildstrubel | N15 | 7.57416 | 46.44123 | 0 | Mariama Diallo, Thomas Bulliard et Stéphanie Morelon | 5 |
| CH_Finges | N16 | 7.56507 | 46.28567 | 4 | Gauvain Demars et Stéphanie Morelon | 5 |
| CH_Pointe_de_Bellevue | N17 | 6.88320 | 46.26294 | 2.5 | Stéphanie Morelon | 5 |
| FR_Pointe_Deux_Pertuis | N18 | 46.23647 | 6.74741 |  | Florian Boucher-Gabriele Casazza-Peter Szovenyi-Elena Conti - Molecular Phylogenetics and Evolution 104 (2016) 60:72 | 1 |
| FR_Sous_Dîne | N19 | 7.24957 | 46.06175 | 0 | Florian Boucher, Marion Deville-Cavellin, Philippe Juillerat et Stéphanie Morelon | 5 |
| FR_Col_des_Aravis | N20 | 45.87011 | 6.4572 |  | Florian Boucher | 8 |
| FR_Grand_Som | N21 | 45.38211 | 5.81102 |  | Camille Voisin | 10 |
| FR_Charmant_Som | N22 | 45.32315 | 5.76232 |  | Florian Boucher | 11 |
| FR_Moucherotte | N23 | 45.14115 | 5.62775 |  | Marion Deville-Cavellin | 10 |
| FR_Pierre_Blanche | N24 | 44.89069 | 5.52131 |  | Florian Boucher | 12 |
| FR_Dévoluy | N25 | 44.68899 | 5.89028 |  | Florian Boucher | 5 |
| DE_Schwarzwald_Höllental | BF | 8.02161 | 47.93773 |  | Florian Boucher-Gabriele Casazza-Peter Szovenyi-Elena Conti - Molecular Phylogenetics and Evolution 104 (2016) 60:72 | 3 |
| DE_Schwarzwald_Höllental | BF | 8.02161 | 47.93773 | 0 | Philippe Juillerat et Stéphanie Morelon | 5 |
| CH_Ramsflue | J1 | 7.99981 | 47.42206 | 2 | Stéphanie Morelon | 5 |
| CH_Geissflue | J2 | 7.80184 | 47.37192 | 3 | Philippe Juillerat et Stéphanie Morelon | 5 |
| CH_Balsthal | J3 | 7.71229 | 47.32678 | 11 | Stéphanie Morelon | 5 |
| CH_Hasenmatt | J4 | 7.45682 | 47.24208 |  | Stéphanie Morelon | 5 |
| CH_Kleinlützel | J5 | 7.412516 | 47.43128 | 4 | Philippe Juillerat et Stéphanie Morelon | 5 |
| CH_Le_Quenet | J6 | 7.3602885 | 47.37959 |  | Philippe Juillerat et Stéphanie Morelon | 2 |
| CH_Court | J7 | 7.34532 | 47.24752 |  | Philippe Juillerat et Stéphanie Morelon | 5 |
| CH_Pichoux | J8 | 7.22780 | 47.28194 | 22 | Philippe Juillerat et Stéphanie Morelon | 5 |
| CH_Chasseral | J9 | 7.02973 | 47.11789 | 1 | Stéphanie Morelon | 5 |
| FR_Roche_Fendue | J10 | 6.76084 | 47.31230 | 9 | Stéphanie Morelon | 2 |
| FR_Roche_de_Chatard | J11 | 6.36849 | 47.34019 | 25 | Stéphanie Morelon | 5 |
| CH_Roches_blanches | J12 | 6.52842 | 46.85893 | 4 | Sergio Rasmann et Stéphanie Morelon | 5 |

**Table S2** DNA concentration (ng/ml) of the purified DNA solutions before PCR amplification, and summary statistics of the ipyrad library assembly of the total sampling of Primula auricula and Primula spectabilis (Dmpx_XSP_1-5)

| **sample** | **DNA concentration**  **(ng/ul)** | | **state** | **Reads raw** | **Reads passed  filter** | **Clusters total** | **Clusters hidepth** | | **Hetero est** | **Error est** | **Reads consens** | **Loci in assembly** | **Clade** | **Geographic_region** |
| --- | --- | --- | --- | --- | --- | --- | --- | --- | --- | --- | --- | --- | --- | --- |
|  | **Before PCR** | **After  PCR** |  |  |  |  |  |  |  |  |  |  |  |  |
| Dmpx_XSP_1 | 3,39 | 13,15 | 7 | 3313980 | 3313339 | 11297 | | 3957 | 0.011586 | 0.000807 | 3431 | 2037 | Outgroup | A |
| Dmpx_XSP_2 | 4,09 | 9,03 | 7 | 5831936 | 5830542 | 20776 | | 4896 | 0.013844 | 0.000841 | 4005 | 2327 | Outgroup | A |
| Dmpx_XSP_3 | 3,52 | 12,41 | 7 | 2047655 | 2047296 | 17076 | | 5172 | 0.012538 | 0.000834 | 4360 | 2512 | Outgroup | A |
| Dmpx_XSP_4 | 6,89 | 8,37 | 7 | 1421121 | 1420876 | 13727 | | 4417 | 0.011678 | 0.000835 | 3801 | 2301 | Outgroup | A |
| Dmpx_XSP_5 | 1,42 | 9,80 | 7 | 1941684 | 1941196 | 16090 | | 4959 | 0.012297 | 0.000803 | 4214 | 2503 | Outgroup | A |
| Dmpx_S1_1 | 5,94 | 10,92 | 7 | 2223336 | 2222548 | 13794 | | 4222 | 0.011309 | 0.001152 | 3724 | 2494 | Clade_1 | A |
| Dmpx_S1_2 | 3,75 | 10,90 | 7 | 2101620 | 2100598 | 12405 | | 4376 | 0.007652 | 0.000993 | 4018 | 2473 | Clade_1 | A |
| Dmpx_S1_3 | 2,82 | 2,98 | 7 | 1896320 | 1895911 | 8555 | | 1995 | 0.025976 | 0.000889 | 1634 | 973 | Clade_1 | A |
| Dmpx_S1_5 | 6,1 | 10,23 | 7 | 2063312 | 2062758 | 10917 | | 4052 | 0.010877 | 0.000929 | 3583 | 2492 | Clade_1 | A |
| Dmpx_S2_1 | 6,04 | 7,22 | 7 | 2408795 | 2408412 | 16781 | | 5545 | 0.010078 | 0.000768 | 4822 | 3085 | Clade_1 | A |
| Dmpx_S2_2 | 6,29 | 8,66 | 7 | 2657280 | 2656916 | 10119 | | 3829 | 0.008725 | 0.000728 | 3396 | 2286 | Clade_1 | A |
| Dmpx_S2_3 | 10,7 | 8,93 | 7 | 1989016 | 1988688 | 14869 | | 4930 | 0.010024 | 0.000758 | 4310 | 2874 | Clade_1 | A |
| Dmpx_S2_4 | 8,32 | 3,77 | 7 | 2927872 | 2927431 | 8925 | | 3318 | 0.010125 | 0.000675 | 2900 | 1942 | Clade_1 | A |
| Dmpx_S3_1 | 3,65 | 9,75 | 7 | 4916298 | 4915515 | 13123 | | 4119 | 0.008644 | 0.000685 | 3660 | 2461 | Clade_1 | A |
| Dmpx_S3_10 | 2,72 | 3,54 | 7 | 3455199 | 3454305 | 16759 | | 3309 | 0.015950 | 0.000944 | 2853 | 1866 | Clade_1 | A |
| Dmpx_S3_2 | 4,79 | 4,95 | 7 | 2057943 | 2057652 | 11758 | | 4100 | 0.008698 | 0.000624 | 3650 | 2693 | Clade_1 | A |
| Dmpx_S3_3 | 1,41 | 8,55 | 7 | 1608207 | 1607919 | 15172 | | 4866 | 0.010198 | 0.000805 | 4266 | 2768 | Clade_1 | A |
| Dmpx_S3_4 | 3,46 | 3,27 | 7 | 205079 | 205041 | 3013 | | 1425 | 0.007854 | 0.001399 | 1306 | 851 | Clade_1 | A |
| Dmpx_S3_6 | 4,55 | 8,30 | 7 | 1800369 | 1799959 | 5883 | | 1478 | 0.008138 | 0.000629 | 1338 | 933 | Clade_1 | A |
| Dmpx_S3_7 | 4,32 | 9,14 | 7 | 1673422 | 1672850 | 9849 | | 2870 | 0.011741 | 0.001068 | 2537 | 1717 | Clade_1 | A |
| Dmpx_S3_9 | 3,61 | 4,65 | 7 | 2400802 | 2400212 | 11013 | | 3811 | 0.012613 | 0.001075 | 3376 | 2348 | Clade_1 | A |
| Dmpx_S4_1 | 2,61 | 10,98 | 7 | 2071398 | 2071063 | 13942 | | 4395 | 0.010362 | 0.000740 | 3843 | 2646 | Clade_1 | A |
| Dmpx_S4_3 | 3,46 | 12,04 | 7 | 2437758 | 2437394 | 14394 | | 4731 | 0.009851 | 0.000755 | 4134 | 2892 | Clade_1 | A |
| Dmpx_S4_4 | 3,36 | 1,06 | 7 | 668943 | 668842 | 4382 | | 2057 | 0.008123 | 0.000860 | 1868 | 1352 | Clade_1 | A |
| Dmpx_S4_5 | 4,13 | 9,82 | 7 | 2151148 | 2150811 | 15040 | | 4889 | 0.010411 | 0.000741 | 4247 | 2891 | Clade_1 | A |
| Dmpx_S5_10 | 5,25 | 14,07 | 7 | 2083076 | 2082447 | 13389 | | 5077 | 0.013075 | 0.001148 | 4434 | 3122 | Clade_1 | A |
| Dmpx_S5_3 | 4,78 | 4,65 | 7 | 1905401 | 1905093 | 16165 | | 5365 | 0.010791 | 0.000827 | 4665 | 2985 | Clade_1 | A |
| Dmpx_S5_4 | 37,4 | 4,17 | 7 | 2298920 | 2298453 | 16698 | | 6246 | 0.008479 | 0.001070 | 5642 | 2860 | Clade_1 | A |
| Dmpx_S5_5 | 14,6 | 3,72 | 7 | 1967342 | 1966962 | 14073 | | 5993 | 0.006861 | 0.001010 | 5517 | 2730 | Clade_1 | A |
| Dmpx_S5_6 | 3,15 | 11,30 | 7 | 1794448 | 1793977 | 11006 | | 4098 | 0.009903 | 0.001146 | 3668 | 2529 | Clade_1 | A |
| Dmpx_S5_7 | 4,63 | 15,23 | 7 | 2225431 | 2224799 | 27240 | | 10786 | 0.015005 | 0.001423 | 9281 | 6596 | Clade_1 | A |
| Dmpx_S5_8 | 4,66 | 12,24 | 7 | 1992807 | 1992136 | 13361 | | 5322 | 0.011235 | 0.001112 | 4794 | 3315 | Clade_1 | A |
| Dmpx_S5_9 | 7,08 | 1,32 | 7 | 3799948 | 3799511 | 9738 | | 2303 | 0.006812 | 0.000675 | 2096 | 1485 | Clade_1 | A |
| Dmpx_S6_10 | 3,64 | 17,37 | 7 | 1822648 | 1822080 | 22364 | | 9927 | 0.010884 | 0.001265 | 8849 | 6060 | Clade_1 | A |
| Dmpx_S6_13 | 5,09 | 12,11 | 7 | 1433298 | 1432929 | 11787 | | 4382 | 0.010217 | 0.001445 | 3892 | 2536 | Clade_1 | A |
| Dmpx_S6_14 | 5,26 | 10,38 | 7 | 2369933 | 2369236 | 13454 | | 3622 | 0.010886 | 0.001138 | 3155 | 2139 | Clade_1 | A |
| Dmpx_S6_6 | 3,81 | 14,86 | 7 | 1721937 | 1721050 | 27659 | | 11680 | 0.009754 | 0.001679 | 10514 | 5972 | Clade_1 | A |
| Dmpx_S6_9 | 2,46 | 3,20 | 7 | 2634173 | 2633471 | 9739 | | 2800 | 0.009764 | 0.000864 | 2516 | 1648 | Clade_1 | A |
| Dmpx_S7_1 | 3,1 | 13,36 | 7 | 2050705 | 2050356 | 9463 | | 3907 | 0.009087 | 0.000822 | 3469 | 2227 | Clade_2 | B |
| Dmpx_S7_2 | 2,02 | 7,82 | 7 | 4116996 | 4116022 | 23909 | | 9596 | 0.007191 | 0.000820 | 8772 | 4355 | Clade_2 | B |
| Dmpx_S7_3 | 1,47 | 7,52 | 7 | 2877771 | 2876395 | 11176 | | 4259 | 0.007289 | 0.000744 | 3851 | 2042 | Clade_2 | B |
| Dmpx_S7_5 | 3,95 | 5,17 | 7 | 4304572 | 4303396 | 35473 | | 13431 | 0.008780 | 0.001005 | 12035 | 5310 | Clade_2 | B |
| Dmpx_S8_1 | 3,76 | 10,05 | 7 | 2456175 | 2455775 | 9893 | | 3988 | 0.009179 | 0.000750 | 3552 | 2333 | Clade_2 | B |
| Dmpx_S8_2 | 10,9 | 1,71 | 7 | 2138606 | 2138250 | 23823 | | 8368 | 0.008008 | 0.001270 | 7464 | 3316 | Clade_2 | B |
| Dmpx_S8_3 | 2,85 | 7,97 | 7 | 2041285 | 2040941 | 15640 | | 5215 | 0.009701 | 0.000733 | 4610 | 3049 | Clade_2 | B |
| Dmpx_S8_4 | 5,17 | 10,82 | 7 | 2624691 | 2624105 | 26502 | | 9713 | 0.007397 | 0.001009 | 8849 | 4664 | Clade_2 | B |
| Dmpx_S8_5 | 5,24 | 9,51 | 7 | 1088104 | 1087868 | 11414 | | 3469 | 0.009588 | 0.001070 | 3105 | 1924 | Clade_2 | B |
| Dmpx_S9_1 | 21,5 | 5,70 | 7 | 1914024 | 1913615 | 13052 | | 5016 | 0.011423 | 0.000979 | 4336 | 2958 | Clade_2 | B |
| Dmpx_S9_2 | 3,03 | 14,95 | 7 | 1460981 | 1460771 | 8775 | | 3974 | 0.009158 | 0.000822 | 3537 | 2472 | Clade_2 | B |
| Dmpx_S9_3 | 1,41 | 14,61 | 7 | 4311991 | 4311035 | 29574 | | 11484 | 0.007325 | 0.000750 | 10566 | 5057 | Clade_2 | B |
| Dmpx_S9_4 | 11,6 | 3,36 | 7 | 2103148 | 2102359 | 11348 | | 4546 | 0.009477 | 0.000815 | 4021 | 2654 | Clade_2 | B |
| Dmpx_S9_5 | 9,16 | 0,99 | 7 | 2755606 | 2755200 | 7518 | | 2361 | 0.007427 | 0.000608 | 2161 | 1412 | Clade_2 | B |
| Dmpx_S10_2 | 3,28 | 13,63 | 7 | 3418929 | 3418200 | 21424 | | 8684 | 0.007022 | 0.000895 | 7975 | 4281 | Clade_2 | B |
| Dmpx_S10_3 | 3,24 | 10,84 | 7 | 1905494 | 1905222 | 8392 | | 3695 | 0.008798 | 0.000726 | 3289 | 2265 | Clade_2 | B |
| Dmpx_S10_4 | 6,04 | 2,90 | 7 | 1760876 | 1760584 | 7859 | | 3181 | 0.008813 | 0.000688 | 2815 | 1974 | Clade_2 | B |
| Dmpx_S10_5 | 4,69 | 8,32 | 7 | 2116322 | 2116007 | 16071 | | 5316 | 0.008261 | 0.000675 | 4768 | 3115 | Clade_2 | B |
| Dmpx_S11_1 | NA | 5,84 | 7 | 1340022 | 1339613 | 11713 | | 4598 | 0.013314 | 0.001314 | 3993 | 2927 | Clade_2 | B |
| Dmpx_S12_2 | 7,48 | 1,63 | 7 | 1887424 | 1887169 | 5704 | | 1364 | 0.007205 | 0.000668 | 1228 | 860 | Clade_2 | B |
| Dmpx_S12_3 | 9,24 | 0,94 | 7 | 1125291 | 1125119 | 4481 | | 1440 | 0.005666 | 0.000859 | 1344 | 908 | Clade_2 | B |
| Dmpx_S12_4 | 5,89 | 6,53 | 7 | 1432786 | 1432573 | 13441 | | 4107 | 0.010930 | 0.000818 | 3539 | 2448 | Clade_2 | B |
| Dmpx_S12_5 | 3,93 | 11,46 | 7 | 526876 | 526765 | 6822 | | 3001 | 0.010298 | 0.001059 | 2682 | 1910 | Clade_2 | B |
| Dmpx_S13_1 | 1,99 | 8,33 | 7 | 2954818 | 2954084 | 10701 | | 4176 | 0.008805 | 0.000703 | 3745 | 2381 | Clade_2 | B |
| Dmpx_S13_2 | 4,9 | 16,09 | 7 | 2454440 | 2453764 | 13389 | | 5298 | 0.009537 | 0.000928 | 4740 | 2322 | Clade_2 | B |
| Dmpx_S13_3 | 2,18 | 4,66 | 7 | 7480069 | 7478721 | 14447 | | 3378 | 0.009479 | 0.000816 | 2967 | 1833 | Clade_2 | B |
| Dmpx_S13_4 | 7,57 | 3,27 | 7 | 2160544 | 2160219 | 8561 | | 3270 | 0.007832 | 0.000737 | 2981 | 2006 | Clade_2 | B |
| Dmpx_S13_5 | 6,08 | 9,47 | 7 | 1636236 | 1635699 | 26060 | | 8748 | 0.005375 | 0.000915 | 8170 | 2822 | Clade_2 | B |
| Dmpx_Ca_1 | NA | 4,69 | 7 | 1523812 | 1523088 | 31412 | | 11024 | 0.007467 | 0.001944 | 10092 | 2632 | Clade_2 | C |
| Dmpx_N1_1 | NA | 4,78 | 7 | 1555889 | 1555122 | 21811 | | 8638 | 0.007824 | 0.001801 | 7871 | 3152 | Clade_2 | D |
| Dmpx_N2_1 | NA | 1,53 | 7 | 2957051 | 2955864 | 17551 | | 5790 | 0.010664 | 0.001274 | 5097 | 2726 | Clade_2 | D |
| Dmpx_N3_1 | 7,17 | 0,77 | 7 | 2671229 | 2670885 | 7186 | | 2380 | 0.005865 | 0.000560 | 2182 | 1561 | Clade_3 | D |
| Dmpx_N3_2 | 4,95 | 15,63 | 7 | 1787417 | 1786442 | 39045 | | 15210 | 0.010615 | 0.001844 | 13757 | 6302 | Clade_3 | D |
| Dmpx_N3_3 | 5,83 | 7,35 | 7 | 1975012 | 1973796 | 16607 | | 5771 | 0.008008 | 0.001279 | 5213 | 2692 | Clade_3 | D |
| Dmpx_N3_5 | 3,15 | 5,16 | 7 | 1051522 | 1051321 | 6468 | | 2466 | 0.007913 | 0.001043 | 2245 | 1557 | Clade_3 | D |
| Dmpx_N4_1 | 7,33 | 1,73 | 7 | 1561164 | 1560951 | 3899 | | 1204 | 0.004340 | 0.000605 | 1119 | 752 | Clade_3 | D |
| Dmpx_N4_3 | 10,2 | 0,84 | 7 | 418549 | 418491 | 3876 | | 1861 | 0.007706 | 0.001113 | 1686 | 1211 | Clade_3 | D |
| Dmpx_N4_4 | 14 | 1,48 | 7 | 1821215 | 1820878 | 9654 | | 4292 | 0.008752 | 0.000773 | 3852 | 2390 | Clade_3 | D |
| Dmpx_N4_5 | 2,07 | 2,24 | 7 | 2409834 | 2409247 | 7512 | | 1863 | 0.010267 | 0.000693 | 1663 | 1131 | Clade_3 | D |
| Dmpx_N5_1 | 7,94 | 1,46 | 7 | 1291892 | 1291709 | 4363 | | 1397 | 0.010082 | 0.000735 | 1251 | 852 | Clade_3 | D |
| Dmpx_N5_2 | 4,11 | 10,83 | 7 | 2040189 | 2039738 | 9275 | | 2775 | 0.009013 | 0.000883 | 2483 | 1680 | Clade_3 | D |
| Dmpx_N5_3 | 3,93 | 2,57 | 7 | 1285898 | 1285440 | 6652 | | 1522 | 0.007006 | 0.001012 | 1410 | 910 | Clade_3 | D |
| Dmpx_N5_4 | 2,65 | 8,69 | 7 | 2814205 | 2812760 | 14297 | | 4796 | 0.008604 | 0.000956 | 4351 | 2867 | Clade_3 | D |
| Dmpx_N5_5 | 3,84 | 9,03 | 7 | 2147195 | 2146660 | 10653 | | 3662 | 0.009794 | 0.000985 | 3262 | 2280 | Clade_3 | D |
| Dmpx_N6_1 | 10 | 0,87 | 7 | 262744 | 262715 | 3222 | | 1463 | 0.006701 | 0.001277 | 1350 | 925 | Clade_3 | E |
| Dmpx_N6_2 | 13,6 | 2,84 | 7 | 2622705 | 2622188 | 14520 | | 5435 | 0.007758 | 0.000923 | 4891 | 2473 | Clade_3 | E |
| Dmpx_N6_3 | 12,4 | 2,90 | 7 | 2548672 | 2548287 | 11342 | | 4584 | 0.007298 | 0.000855 | 4182 | 2052 | Clade_3 | E |
| Dmpx_N6_4 | 23,3 | 3,05 | 7 | 2667211 | 2666747 | 13477 | | 5189 | 0.007538 | 0.000861 | 4690 | 2482 | Clade_3 | E |
| Dmpx_N6_5 | 18,8 | 3,31 | 7 | 1901119 | 1900716 | 15416 | | 5860 | 0.007328 | 0.001066 | 5327 | 2546 | Clade_3 | E |
| Dmpx_N7_1 | NA | 1,93 | 7 | 1084386 | 1084047 | 9403 | | 4843 | 0.006105 | 0.001093 | 4510 | 1866 | Clade_3 | E |
| Dmpx_N8_1 | 10,2 | 1,32 | 7 | 1526145 | 1525914 | 7886 | | 3178 | 0.007293 | 0.000815 | 2882 | 1984 | Clade_3 | E |
| Dmpx_N8_2 | 11,7 | 3,74 | 7 | 1423188 | 1422840 | 15970 | | 5751 | 0.007497 | 0.001241 | 5143 | 2345 | Clade_3 | E |
| Dmpx_N8_4 | 17,8 | 1,83 | 7 | 454834 | 454355 | 11299 | | 3144 | 0.008990 | 0.001046 | 2836 | 1904 | Clade_3 | E |
| Dmpx_N8_5 | 13,4 | 1,74 | 7 | 1970224 | 1969945 | 10773 | | 4579 | 0.007151 | 0.000811 | 4163 | 2256 | Clade_3 | E |
| Dmpx_N9_1 | 4,8 | 14,38 | 7 | 2100561 | 2099737 | 16304 | | 5808 | 0.008361 | 0.001497 | 5260 | 2619 | Clade_3 | E |
| Dmpx_N9_3 | 3,43 | 4,38 | 7 | 1982094 | 1981177 | 21511 | | 7765 | 0.008277 | 0.001702 | 7060 | 2492 | Clade_3 | E |
| Dmpx_N9_4 | 5,86 | 14,81 | 7 | 1759318 | 1758711 | 30572 | | 11291 | 0.012156 | 0.001822 | 10017 | 6537 | Clade_3 | E |
| Dmpx_N9_5 | 6,55 | 17,63 | 7 | 2185770 | 2184678 | 30123 | | 12305 | 0.012773 | 0.001454 | 10928 | 7212 | Clade_3 | E |
| Dmpx_N10_1 | NA | 2,97 | 7 | 2066033 | 2065652 | 14412 | | 6388 | 0.006433 | 0.000981 | 5885 | 3241 | Clade_3 | E |
| Dmpx_N11_1 | 17,9 | 3,15 | 7 | 1807840 | 1807464 | 10479 | | 3721 | 0.010829 | 0.000968 | 3195 | 2125 | Clade_3 | E |
| Dmpx_N11_2 | 6,5 | 16,45 | 7 | 1934878 | 1933939 | 32253 | | 12452 | 0.013138 | 0.001506 | 11002 | 7198 | Clade_3 | E |
| Dmpx_N11_4 | 11 | 6,12 | 7 | 2397857 | 2397520 | 24338 | | 6630 | 0.014469 | 0.001184 | 5552 | 3319 | Clade_3 | E |
| Dmpx_N12_1 | 6,59 | 11,30 | 7 | 2099924 | 2099245 | 16111 | | 5821 | 0.010219 | 0.001380 | 5269 | 3177 | Clade_3 | E |
| Dmpx_N12_2 | 11,4 | 5,55 | 7 | 2106895 | 2106497 | 17494 | | 6167 | 0.008407 | 0.000844 | 5549 | 3558 | Clade_3 | E |
| Dmpx_N12_3 | 3,28 | 3,48 | 7 | 1315862 | 1315521 | 6775 | | 1508 | 0.014719 | 0.001031 | 1337 | 876 | Clade_3 | E |
| Dmpx_N12_4 | 5,54 | 18,28 | 7 | 2174500 | 2173623 | 32833 | | 12330 | 0.010955 | 0.001712 | 11080 | 6515 | Clade_3 | E |
| Dmpx_N12_5 | 9,84 | 1,13 | 7 | 614059 | 613964 | 4441 | | 1959 | 0.005825 | 0.000966 | 1820 | 1229 | Clade_3 | E |
| Dmpx_N13_1 | 12,5 | 1,31 | 7 | 1856812 | 1856532 | 11945 | | 4803 | 0.007188 | 0.000914 | 4370 | 2433 | Clade_3 | E |
| Dmpx_N13_2 | 13 | 2,09 | 7 | 2084886 | 2084499 | 13533 | | 4933 | 0.008606 | 0.000868 | 4341 | 2402 | Clade_3 | E |
| Dmpx_N13_3 | 20,5 | 3,23 | 7 | 2990939 | 2990308 | 18191 | | 7729 | 0.005837 | 0.000949 | 7180 | 2791 | Clade_3 | E |
| Dmpx_N13_5 | 8,73 | 1,12 | 7 | 1575755 | 1575304 | 9274 | | 3141 | 0.005521 | 0.000870 | 2934 | 1765 | Clade_3 | E |
| Dmpx_N14_1 | 4,4 | 8,73 | 7 | 2659700 | 2659089 | 14480 | | 4635 | 0.009960 | 0.001129 | 4128 | 2395 | Clade_3 | E |
| Dmpx_N14_2 | 3,41 | 6,74 | 7 | 2419575 | 2418987 | 17918 | | 4352 | 0.010433 | 0.001481 | 3834 | 1976 | Clade_3 | E |
| Dmpx_N14_3 | 4,57 | 14,68 | 7 | 2438281 | 2437526 | 15703 | | 4914 | 0.011897 | 0.001279 | 4370 | 2925 | Clade_3 | E |
| Dmpx_N14_4 | 4,2 | 12,69 | 7 | 1792035 | 1791434 | 13730 | | 4768 | 0.010499 | 0.001405 | 4299 | 2779 | Clade_3 | E |
| Dmpx_N15_1 | 7 | 2,61 | 7 | 2463603 | 2463171 | 11653 | | 4016 | 0.009748 | 0.000906 | 3543 | 2225 | Clade_3 | E |
| Dmpx_N15_2 | 2,96 | 0,00 | 7 | 587070 | 586990 | 3545 | | 1354 | 0.004441 | 0.000976 | 1276 | 869 | Clade_3 | E |
| Dmpx_N15_3 | 1,89 | 3,14 | 7 | 978363 | 978194 | 7017 | | 3278 | 0.005873 | 0.000716 | 3090 | 1660 | Clade_3 | E |
| Dmpx_N15_4 | 1,89 | 13,36 | 7 | 2294810 | 2294488 | 8001 | | 2935 | 0.008442 | 0.000652 | 2619 | 1901 | Clade_3 | E |
| Dmpx_N15_5 | 1,82 | 3,04 | 7 | 2440980 | 2440569 | 14194 | | 5616 | 0.009353 | 0.000912 | 5003 | 2569 | Clade_3 | E |
| Dmpx_N16_1 | 8,22 | 2,51 | 7 | 2756674 | 2756297 | 12757 | | 3985 | 0.006042 | 0.000838 | 3635 | 2067 | Clade_3 | E |
| Dmpx_N16_2 | 2,72 | 6,05 | 7 | 3566168 | 3565212 | 16942 | | 5150 | 0.009692 | 0.000999 | 4636 | 2774 | Clade_3 | E |
| Dmpx_N16_3 | 6,24 | 5,16 | 7 | 2303398 | 2302686 | 13288 | | 4782 | 0.008646 | 0.001115 | 4338 | 2890 | Clade_3 | E |
| Dmpx_N16_5 | 5,62 | 8,26 | 7 | 2119803 | 2119194 | 15457 | | 6061 | 0.009161 | 0.001142 | 5463 | 3276 | Clade_3 | E |
| Dmpx_N17_1 | 10,7 | 1,11 | 7 | 282569 | 282531 | 2661 | | 1227 | 0.003617 | 0.001105 | 1170 | 657 | Clade_3 | E |
| Dmpx_N17_2 | 6,82 | 12,26 | 7 | 1761320 | 1760520 | 23400 | | 9076 | 0.008287 | 0.001667 | 8254 | 3301 | Clade_3 | E |
| Dmpx_N17_3 | 15,6 | 3,25 | 7 | 973213 | 973078 | 3529 | | 1550 | 0.005325 | 0.000659 | 1454 | 903 | Clade_3 | E |
| Dmpx_N17_4 | 5,8 | 10,55 | 7 | 2120431 | 2119781 | 15079 | | 5550 | 0.009572 | 0.001269 | 5004 | 2911 | Clade_3 | E |
| Dmpx_N17_5 | 4,59 | 18,61 | 7 | 2488653 | 2487779 | 27786 | | 10858 | 0.011913 | 0.001519 | 9717 | 6649 | Clade_3 | E |
| Dmpx_N18_1 | NA | 2,19 | 7 | 2881293 | 2880342 | 20157 | | 6065 | 0.013353 | 0.001178 | 5330 | 3756 | Clade_3 | F |
| Dmpx_N19_1 | 8 | 1,11 | 7 | 267086 | 267052 | 2060 | | 838 | 0.002973 | 0.000917 | 804 | 531 | Clade_3 | F |
| Dmpx_N19_2 | 9,24 | 1,17 | 7 | 511113 | 511051 | 2366 | | 1021 | 0.003131 | 0.000709 | 969 | 672 | Clade_3 | F |
| Dmpx_N19_4 | 11,8 | 3,00 | 7 | 2219164 | 2218680 | 13024 | | 4795 | 0.009740 | 0.000913 | 4248 | 2407 | Clade_3 | F |
| Dmpx_N19_5 | 8,3 | 1,56 | 7 | 1966739 | 1966435 | 6240 | | 2407 | 0.005719 | 0.000656 | 2215 | 1497 | Clade_3 | F |
| Dmpx_N20_1 | 4,72 | 10,04 | 7 | 1172292 | 1171846 | 9670 | | 3243 | 0.010570 | 0.001278 | 2883 | 1801 | Clade_3 | F |
| Dmpx_N20_2 | 3,88 | 10,60 | 7 | 1475137 | 1474379 | 8766 | | 2735 | 0.019103 | 0.001044 | 2330 | 1503 | Clade_3 | F |
| Dmpx_N20_4 | 3,77 | 12,84 | 7 | 1939424 | 1938762 | 13426 | | 4497 | 0.011926 | 0.001293 | 4062 | 2580 | Clade_3 | F |
| Dmpx_N20_5 | 2,73 | 3,71 | 7 | 2183677 | 2183104 | 10168 | | 3429 | 0.009733 | 0.000816 | 3121 | 2033 | Clade_3 | F |
| Dmpx_N20_6 | 2,87 | 5,75 | 7 | 1659967 | 1659510 | 6780 | | 1809 | 0.011208 | 0.000937 | 1604 | 1103 | Clade_3 | F |
| Dmpx_N20_8 | 4,76 | 7,00 | 7 | 2123220 | 2122423 | 13775 | | 3953 | 0.008360 | 0.001122 | 3580 | 1681 | Clade_3 | F |
| Dmpx_N21_10 | 4,74 | 5,49 | 7 | 824525 | 824289 | 5839 | | 2357 | 0.008029 | 0.001144 | 2148 | 1511 | Clade_3 | F |
| Dmpx_N21_2 | 4,99 | 13,58 | 7 | 1956587 | 1956068 | 12801 | | 5092 | 0.011869 | 0.001094 | 4503 | 3268 | Clade_3 | F |
| Dmpx_N21_3 | 5,72 | 14,48 | 7 | 2324458 | 2323547 | 24295 | | 8563 | 0.012468 | 0.001633 | 7603 | 3978 | Clade_3 | F |
| Dmpx_N21_4 | 5,19 | 8,51 | 7 | 1982605 | 1982091 | 13201 | | 4790 | 0.009371 | 0.001142 | 4269 | 2619 | Clade_3 | F |
| Dmpx_N21_5 | 6,36 | 3,69 | 7 | 2384389 | 2383744 | 12661 | | 3788 | 0.012346 | 0.001049 | 3290 | 2390 | Clade_3 | F |
| Dmpx_N21_6 | 10,3 | 0,97 | 7 | 757452 | 757338 | 5459 | | 2203 | 0.006552 | 0.001040 | 2033 | 1472 | Clade_3 | F |
| Dmpx_N21_7 | 10,1 | 13,46 | 7 | 1300059 | 1299730 | 8127 | | 3379 | 0.008902 | 0.000887 | 3036 | 2218 | Clade_3 | F |
| Dmpx_N22_1 | 4,05 | 18,00 | 7 | 2125805 | 2125225 | 25974 | | 10376 | 0.014685 | 0.001516 | 8978 | 7010 | Clade_3 | F |
| Dmpx_N22_10 | 4,46 | 20,44 | 7 | 2091548 | 2090748 | 22791 | | 10511 | 0.010970 | 0.001182 | 9483 | 6908 | Clade_3 | F |
| Dmpx_N22_4 | 4,91 | 16,95 | 7 | 1849585 | 1848970 | 20499 | | 9904 | 0.011628 | 0.001107 | 8857 | 7028 | Clade_3 | F |
| Dmpx_N22_7 | 4,5 | 15,20 | 7 | 1885505 | 1884942 | 19208 | | 8357 | 0.009182 | 0.001409 | 7592 | 5744 | Clade_3 | F |
| Dmpx_N22_8 | 5,39 | 13,83 | 7 | 1686891 | 1686393 | 9183 | | 4154 | 0.009632 | 0.000987 | 3795 | 2702 | Clade_3 | F |
| Dmpx_N22_9 | 6,13 | 16,81 | 7 | 1651277 | 1650658 | 20101 | | 9624 | 0.010478 | 0.001246 | 8604 | 6549 | Clade_3 | F |
| Dmpx_N23_1 | 20,5 | 7,91 | 7 | 1564754 | 1564151 | 8938 | | 3116 | 0.006997 | 0.000918 | 2857 | 1869 | Clade_3 | F |
| Dmpx_N23_10 | 2,96 | 4,45 | 7 | 2242438 | 2241898 | 8697 | | 2537 | 0.008674 | 0.000838 | 2284 | 1608 | Clade_3 | F |
| Dmpx_N23_3 | 19,6 | 10,99 | 7 | 1657681 | 1657287 | 8632 | | 3723 | 0.008590 | 0.000884 | 3414 | 2457 | Clade_3 | F |
| Dmpx_N23_5 | 22,8 | 3,79 | 7 | 1479624 | 1479306 | 7677 | | 2258 | 0.010234 | 0.001185 | 2031 | 1451 | Clade_3 | F |
| Dmpx_N23_6 | 38,1 | 5,74 | 7 | 1533322 | 1532984 | 7461 | | 2680 | 0.007002 | 0.000903 | 2431 | 1743 | Clade_3 | F |
| Dmpx_N23_7 | 8,19 | 9,67 | 7 | 1417937 | 1417373 | 6612 | | 2209 | 0.007245 | 0.000781 | 2013 | 1411 | Clade_3 | F |
| Dmpx_N23_8 | 13,9 | 18,51 | 7 | 1907743 | 1907093 | 21344 | | 9185 | 0.011708 | 0.001451 | 8187 | 6507 | Clade_3 | F |
| Dmpx_N23_9 | 14,7 | 28,87 | 7 | 1856024 | 1855171 | 22056 | | 9125 | 0.014057 | 0.001351 | 8048 | 6570 | Clade_3 | F |
| Dmpx_N24_1 | 3,02 | 8,36 | 7 | 1934441 | 1933793 | 9790 | | 2935 | 0.009160 | 0.001007 | 2646 | 1614 | Clade_3 | F |
| Dmpx_N24_10 | 4,57 | 22,09 | 7 | 1906121 | 1905083 | 22684 | | 9844 | 0.010306 | 0.001502 | 8864 | 6479 | Clade_3 | F |
| Dmpx_N24_11 | 6,3 | 22,34 | 7 | 1937472 | 1936538 | 24183 | | 10849 | 0.009479 | 0.001229 | 9872 | 6828 | Clade_3 | F |
| Dmpx_N24_12 | 4,75 | 20,59 | 7 | 2419969 | 2419121 | 27139 | | 11797 | 0.012642 | 0.001370 | 10467 | 7632 | Clade_3 | F |
| Dmpx_N24_5 | 3,82 | 14,26 | 7 | 1727053 | 1726602 | 12803 | | 5688 | 0.008849 | 0.001196 | 5198 | 3030 | Clade_3 | F |
| Dmpx_N24_7 | 6,3 | 12,60 | 7 | 1884368 | 1883891 | 11705 | | 4174 | 0.008020 | 0.001159 | 3809 | 2556 | Clade_3 | F |
| Dmpx_N24_8 | 3,84 | 14,73 | 7 | 2039471 | 2038965 | 11909 | | 4116 | 0.010763 | 0.001167 | 3699 | 2563 | Clade_3 | F |
| Dmpx_N24_9 | 17,4 | 11,06 | 7 | 1385276 | 1384906 | 7707 | | 3244 | 0.008624 | 0.001020 | 2934 | 2090 | Clade_3 | F |
| Dmpx_N25_1 | 1,02 | 2,49 | 7 | 2591824 | 2591428 | 11933 | | 5227 | 0.007588 | 0.000863 | 4727 | 2321 | Clade_3 | F |
| Dmpx_N25_2 | 4,38 | 2,82 | 7 | 1965082 | 1964183 | 25629 | | 9482 | 0.011090 | 0.001661 | 8339 | 3082 | Clade_3 | F |
| Dmpx_N25_3 | 8,29 | 3,69 | 7 | 1284561 | 1284373 | 11119 | | 4675 | 0.009396 | 0.000941 | 4218 | 2645 | Clade_3 | F |
| Dmpx_N25_4 | 5,82 | 2,52 | 7 | 1602897 | 1602090 | 15938 | | 6361 | 0.009282 | 0.001404 | 5719 | 3058 | Clade_3 | F |
| Dmpx_N25_5 | 1,92 | 2,99 | 7 | 2921279 | 2920759 | 25172 | | 10838 | 0.009144 | 0.001317 | 9663 | 2907 | Clade_3 | F |
| Dmpx_BF_1 | 6,19 | 13,03 | 7 | 2382951 | 2382265 | 14126 | | 5345 | 0.010972 | 0.001152 | 4796 | 3058 | Clade_3 | G |
| Dmpx_BF_2 | 3,19 | 6,03 | 7 | 1778030 | 1777622 | 7014 | | 2121 | 0.010984 | 0.000740 | 1904 | 1277 | Clade_3 | G |
| Dmpx_BF_3 | 5,58 | 7,51 | 7 | 2781210 | 2780426 | 18403 | | 5018 | 0.011551 | 0.001291 | 4394 | 2697 | Clade_3 | G |
| Dmpx_BF_4 | 6,19 | 13,16 | 7 | 2290365 | 2289773 | 13402 | | 4952 | 0.012836 | 0.001238 | 4397 | 3201 | Clade_3 | G |
| Dmpx_BF_5 | 5,9 | 7,28 | 7 | 2045913 | 2045142 | 14064 | | 5280 | 0.010599 | 0.001187 | 4678 | 2977 | Clade_3 | G |
| Dmpx_BF_6 | NA | 2,57 | 7 | 3005988 | 3004973 | 34805 | | 14871 | 0.006402 | 0.001311 | 13725 | 3366 | Clade_3 | G |
| Dmpx_BF_7 | NA | 1,10 | 7 | 1336091 | 1334785 | 26233 | | 10771 | 0.008233 | 0.002200 | 9819 | 3690 | Clade_3 | G |
| Dmpx_BF_8 | NA | 4,97 | 7 | 1928942 | 1928630 | 14302 | | 6727 | 0.007182 | 0.001071 | 6193 | 2947 | Clade_3 | G |
| Dmpx_J1_1 | 5,39 | 1,48 | 7 | 2447557 | 2447108 | 14730 | | 5460 | 0.006740 | 0.000998 | 4982 | 2558 | Clade_3 | H |
| Dmpx_J1_2 | 5,05 | 9,82 | 7 | 2203220 | 2202358 | 11717 | | 3611 | 0.007602 | 0.000954 | 3299 | 2052 | Clade_3 | H |
| Dmpx_J1_3 | 3,98 | 12,80 | 7 | 2264014 | 2263404 | 12177 | | 4671 | 0.011937 | 0.001055 | 4107 | 2972 | Clade_3 | H |
| Dmpx_J1_4 | 6,68 | 7,41 | 7 | 2393696 | 2393158 | 12214 | | 4466 | 0.009191 | 0.000944 | 4004 | 2589 | Clade_3 | H |
| Dmpx_J1_5 | 5,61 | 21,05 | 7 | 2044555 | 2043543 | 31967 | | 14496 | 0.009696 | 0.001682 | 13153 | 7594 | Clade_3 | H |
| Dmpx_J2_1 | 3,97 | 15,62 | 7 | 1574354 | 1573622 | 15258 | | 6907 | 0.008091 | 0.001264 | 6384 | 4423 | Clade_3 | H |
| Dmpx_J2_2 | 5,32 | 33,21 | 7 | 1973805 | 1972674 | 25280 | | 11209 | 0.013651 | 0.001442 | 9896 | 7325 | Clade_3 | H |
| Dmpx_J2_3 | 2,49 | 14,42 | 7 | 1922844 | 1922279 | 9161 | | 3562 | 0.010705 | 0.000912 | 3232 | 2320 | Clade_3 | H |
| Dmpx_J2_4 | 3,28 | 6,00 | 7 | 2219470 | 2218979 | 12710 | | 4258 | 0.009734 | 0.000978 | 3789 | 2314 | Clade_3 | H |
| Dmpx_J2_5 | 4,34 | 9,79 | 7 | 3107424 | 3106083 | 15719 | | 5331 | 0.012215 | 0.001104 | 4707 | 2834 | Clade_3 | H |
| Dmpx_J3_2 | 4,31 | 6,79 | 7 | 2135814 | 2135513 | 13380 | | 4575 | 0.010446 | 0.000714 | 3993 | 2839 | Clade_3 | H |
| Dmpx_J3_3 | 2,33 | 8,77 | 7 | 6200673 | 6199567 | 22099 | | 5713 | 0.010153 | 0.000773 | 5021 | 2767 | Clade_3 | H |
| Dmpx_J3_4 | 6,02 | 2,50 | 7 | 2361953 | 2361583 | 7588 | | 2052 | 0.005805 | 0.000740 | 1878 | 1297 | Clade_3 | H |
| Dmpx_J3_5 | 3,22 | 15,85 | 7 | 2222326 | 2221956 | 16599 | | 5461 | 0.010463 | 0.000764 | 4814 | 3032 | Clade_3 | H |
| Dmpx_J4_1 | 6,2 | 12,94 | 7 | 1866606 | 1865641 | 21756 | | 8461 | 0.007685 | 0.001787 | 7731 | 3168 | Clade_3 | H |
| Dmpx_J4_2 | 5,62 | 15,29 | 7 | 2157195 | 2155665 | 28876 | | 11908 | 0.010716 | 0.001343 | 10723 | 6571 | Clade_3 | H |
| Dmpx_J4_3 | 2,98 | 6,61 | 7 | 1666430 | 1665952 | 9256 | | 3609 | 0.008752 | 0.001065 | 3267 | 1986 | Clade_3 | H |
| Dmpx_J4_4 | 3,22 | 4,17 | 7 | 1546819 | 1546507 | 5157 | | 1323 | 0.005597 | 0.000788 | 1239 | 780 | Clade_3 | H |
| Dmpx_J4_5 | 5,59 | 12,53 | 7 | 2077727 | 2076313 | 29501 | | 10703 | 0.009192 | 0.002058 | 9619 | 3643 | Clade_3 | H |
| Dmpx_J5_2 | 1,86 | 11,20 | 7 | 1655051 | 1654571 | 8939 | | 3863 | 0.008166 | 0.000914 | 3545 | 2540 | Clade_3 | H |
| Dmpx_J5_3 | 5,25 | 6,04 | 7 | 2460432 | 2459814 | 13478 | | 4348 | 0.011023 | 0.000901 | 3828 | 2615 | Clade_3 | H |
| Dmpx_J5_4 | 5,66 | 10,44 | 7 | 2398065 | 2397394 | 14576 | | 4398 | 0.013331 | 0.001013 | 3816 | 2777 | Clade_3 | H |
| Dmpx_J5_5 | 3,7 | 4,21 | 7 | 1305083 | 1304803 | 5914 | | 2132 | 0.005408 | 0.000821 | 1992 | 1434 | Clade_3 | H |
| Dmpx_J6_1 | 3,24 | 1,86 | 7 | 1636188 | 1635923 | 14675 | | 4771 | 0.011212 | 0.001148 | 4168 | 2480 | Clade_3 | H |
| Dmpx_J6_2 | 1,36 | 19,35 | 7 | 1633636 | 1633052 | 21808 | | 9772 | 0.011762 | 0.001538 | 8627 | 6693 | Clade_3 | H |
| Dmpx_J7_1 | 0,265 | 7,75 | 7 | 2188410 | 2188056 | 16410 | | 5612 | 0.010641 | 0.000660 | 4888 | 3376 | Clade_3 | H |
| Dmpx_J7_2 | 1,16 | 17,83 | 7 | 2439132 | 2438714 | 19629 | | 6104 | 0.010371 | 0.000867 | 5318 | 3238 | Clade_3 | H |
| Dmpx_J7_3 | 9,5 | 8,79 | 7 | 1848426 | 1848106 | 14580 | | 4514 | 0.010897 | 0.000836 | 3947 | 2789 | Clade_3 | H |
| Dmpx_J7_4 | 4,5 | 15,21 | 7 | 1595806 | 1595560 | 13829 | | 4531 | 0.010895 | 0.000738 | 3952 | 2786 | Clade_3 | H |
| Dmpx_J7_5 | 1,85 | 10,06 | 7 | 1606606 | 1606349 | 15679 | | 5114 | 0.010750 | 0.000849 | 4479 | 2955 | Clade_3 | H |
| Dmpx_J8_1 | 3,37 | 8,65 | 7 | 1987551 | 1987184 | 9619 | | 3212 | 0.009176 | 0.000878 | 2862 | 2041 | Clade_3 | H |
| Dmpx_J8_2 | 4,72 | 2,10 | 7 | 2214924 | 2214645 | 8387 | | 2482 | 0.007684 | 0.000714 | 2238 | 1595 | Clade_3 | H |
| Dmpx_J8_3 | 4,72 | 0,70 | 7 | 3816905 | 3816387 | 7623 | | 2166 | 0.005965 | 0.000598 | 1960 | 1419 | Clade_3 | H |
| Dmpx_J8_4 | 4,42 | 17,69 | 7 | 2452733 | 2451353 | 39313 | | 14498 | 0.011800 | 0.001859 | 12915 | 6904 | Clade_3 | H |
| Dmpx_J8_5 | 3,3 | 10,64 | 7 | 2068905 | 2068362 | 12948 | | 4942 | 0.010204 | 0.001059 | 4403 | 2850 | Clade_3 | H |
| Dmpx_J9_1 | 3,19 | 8,78 | 7 | 1598725 | 1598336 | 10369 | | 3159 | 0.011161 | 0.001235 | 2802 | 1919 | Clade_3 | H |
| Dmpx_J9_10 | 3,5 | 1,01 | 7 | 7815294 | 7814207 | 21479 | | 4671 | 0.014896 | 0.000651 | 3967 | 2885 | Clade_3 | H |
| Dmpx_J9_3 | 1,58 | 14,83 | 7 | 1970137 | 1969498 | 23930 | | 9380 | 0.010264 | 0.001613 | 8444 | 5446 | Clade_3 | H |
| Dmpx_J9_4 | 3,79 | 9,47 | 7 | 1980266 | 1979789 | 10102 | | 3236 | 0.006891 | 0.000983 | 2989 | 1800 | Clade_3 | H |
| Dmpx_J9_5 | 3,5 | 10,37 | 7 | 2014187 | 2013123 | 11632 | | 3579 | 0.009327 | 0.001019 | 3277 | 1726 | Clade_3 | H |
| Dmpx_J9_6 | 3,19 | 15,50 | 7 | 1871225 | 1870555 | 22756 | | 9628 | 0.013789 | 0.001401 | 8403 | 6453 | Clade_3 | H |
| Dmpx_J9_7_r2 | 8,05 | 29,40 | 7 | 2063092 | 2061779 | 23950 | | 10625 | 0.013912 | 0.001499 | 9377 | 6996 | Clade_3 | H |
| Dmpx_J10_1 | 5,62 | 3,09 | 7 | 1772203 | 1771847 | 4726 | | 1079 | 0.003466 | 0.000618 | 1017 | 613 | Clade_3 | H |
| Dmpx_J10_2 | 4,29 | 3,38 | 7 | 1881723 | 1881271 | 6547 | | 1955 | 0.014920 | 0.000760 | 1728 | 1127 | Clade_3 | H |
| Dmpx_J11_1 | 4,27 | 18,31 | 7 | 2207287 | 2206358 | 25016 | | 9269 | 0.014319 | 0.001516 | 8072 | 6129 | Clade_3 | H |
| Dmpx_J11_2 | 4,48 | 4,74 | 7 | 2020975 | 2020616 | 9450 | | 3711 | 0.009181 | 0.000780 | 3310 | 2238 | Clade_3 | H |
| Dmpx_J11_3 | 4,21 | 14,37 | 7 | 2328138 | 2327556 | 12698 | | 5097 | 0.011000 | 0.001150 | 4586 | 3230 | Clade_3 | H |
| Dmpx_J11_4 | 5,79 | 2,47 | 7 | 2277262 | 2276949 | 7239 | | 2050 | 0.009476 | 0.000857 | 1819 | 1286 | Clade_3 | H |
| Dmpx_J11_5 | 6,52 | 2,98 | 7 | 1060627 | 1060394 | 4606 | | 1162 | 0.004422 | 0.000834 | 1088 | 744 | Clade_3 | H |
| Dmpx_J12_1 | 4,94 | 5,90 | 7 | 2346564 | 2345883 | 7888 | | 2100 | 0.006781 | 0.000749 | 1929 | 1257 | Clade_3 | H |
| Dmpx_J12_2 | 4,23 | 4,04 | 7 | 2243257 | 2242665 | 9654 | | 2985 | 0.010564 | 0.000823 | 2677 | 1747 | Clade_3 | H |
| Dmpx_J12_3 | 3,8 | 9,47 | 7 | 3522110 | 3520394 | 31327 | | 9791 | 0.008732 | 0.001596 | 8818 | 2955 | Clade_3 | H |
| Dmpx_J12_4 | 6,61 | 4,18 | 7 | 1746874 | 1746526 | 13342 | | 5706 | 0.008062 | 0.000850 | 5201 | 3377 | Clade_3 | H |
| Dmpx_J12_5 | 2,53 | 3,81 | 7 | 2034151 | 2033274 | 32561 | | 11071 | 0.008509 | 0.001393 | 10021 | 3789 | Clade_3 | H |

**Table S3** *Primula auricula* and *Primula spectabilis* demultiplexed sequences: NCBI availability

| **Name** | **Accession** | **Corresponding URLs** |
| --- | --- | --- |
| Dmpx_BF_1 | SAMN47739274 | https://www.ncbi.nlm.nih.gov/sra/47739274 |
| Dmpx_BF_2 | SAMN47739275 | https://www.ncbi.nlm.nih.gov/sra/47739275 |
| Dmpx_BF_3 | SAMN47739276 | https://www.ncbi.nlm.nih.gov/sra/47739276 |
| Dmpx_BF_4 | SAMN47739277 | https://www.ncbi.nlm.nih.gov/sra/47739277 |
| Dmpx_BF_5 | SAMN47739278 | https://www.ncbi.nlm.nih.gov/sra/47739278 |
| Dmpx_BF_6 | SAMN47739279 | https://www.ncbi.nlm.nih.gov/sra/47739279 |
| Dmpx_BF_7 | SAMN47739280 | https://www.ncbi.nlm.nih.gov/sra/47739280 |
| Dmpx_BF_8 | SAMN47739281 | https://www.ncbi.nlm.nih.gov/sra/47739281 |
| Dmpx_Ca_1 | SAMN47739282 | https://www.ncbi.nlm.nih.gov/sra/47739282 |
| Dmpx_J10_1 | SAMN47739283 | https://www.ncbi.nlm.nih.gov/sra/47739283 |
| Dmpx_J10_2 | SAMN47739284 | https://www.ncbi.nlm.nih.gov/sra/47739284 |
| Dmpx_J11_1 | SAMN47739285 | https://www.ncbi.nlm.nih.gov/sra/47739285 |
| Dmpx_J11_2 | SAMN47739286 | https://www.ncbi.nlm.nih.gov/sra/47739286 |
| Dmpx_J11_3 | SAMN47739287 | https://www.ncbi.nlm.nih.gov/sra/47739287 |
| Dmpx_J11_4 | SAMN47739288 | https://www.ncbi.nlm.nih.gov/sra/47739288 |
| Dmpx_J11_5 | SAMN47739289 | https://www.ncbi.nlm.nih.gov/sra/47739289 |
| Dmpx_J1_1 | SAMN47739290 | https://www.ncbi.nlm.nih.gov/sra/47739290 |
| Dmpx_J12_1 | SAMN47739291 | https://www.ncbi.nlm.nih.gov/sra/47739291 |
| Dmpx_J12_2 | SAMN47739292 | https://www.ncbi.nlm.nih.gov/sra/47739292 |
| Dmpx_J12_3 | SAMN47739293 | https://www.ncbi.nlm.nih.gov/sra/47739293 |
| Dmpx_J12_4 | SAMN47739294 | https://www.ncbi.nlm.nih.gov/sra/47739294 |
| Dmpx_J12_5 | SAMN47739295 | https://www.ncbi.nlm.nih.gov/sra/47739295 |
| Dmpx_J1_2 | SAMN47739296 | https://www.ncbi.nlm.nih.gov/sra/47739296 |
| Dmpx_J1_3 | SAMN47739297 | https://www.ncbi.nlm.nih.gov/sra/47739297 |
| Dmpx_J1_4 | SAMN47739298 | https://www.ncbi.nlm.nih.gov/sra/47739298 |
| Dmpx_J1_5 | SAMN47739299 | https://www.ncbi.nlm.nih.gov/sra/47739299 |
| Dmpx_J2_1 | SAMN47739300 | https://www.ncbi.nlm.nih.gov/sra/47739300 |
| Dmpx_J2_2 | SAMN47739301 | https://www.ncbi.nlm.nih.gov/sra/47739301 |
| Dmpx_J2_3 | SAMN47739302 | https://www.ncbi.nlm.nih.gov/sra/47739302 |
| Dmpx_J2_4 | SAMN47739303 | https://www.ncbi.nlm.nih.gov/sra/47739303 |
| Dmpx_J2_5 | SAMN47739304 | https://www.ncbi.nlm.nih.gov/sra/47739304 |
| Dmpx_J3_1 | SAMN47739305 | https://www.ncbi.nlm.nih.gov/sra/47739305 |
| Dmpx_J3_2 | SAMN47739306 | https://www.ncbi.nlm.nih.gov/sra/47739306 |
| Dmpx_J3_3 | SAMN47739307 | https://www.ncbi.nlm.nih.gov/sra/47739307 |
| Dmpx_J3_4 | SAMN47739308 | https://www.ncbi.nlm.nih.gov/sra/47739308 |
| Dmpx_J3_5 | SAMN47739309 | https://www.ncbi.nlm.nih.gov/sra/47739309 |
| Dmpx_J4_1 | SAMN47739310 | https://www.ncbi.nlm.nih.gov/sra/47739310 |
| Dmpx_J4_2 | SAMN47739311 | https://www.ncbi.nlm.nih.gov/sra/47739311 |
| Dmpx_J4_3 | SAMN47739312 | https://www.ncbi.nlm.nih.gov/sra/47739312 |
| Dmpx_J4_4 | SAMN47739313 | https://www.ncbi.nlm.nih.gov/sra/47739313 |
| Dmpx_J4_5 | SAMN47739314 | https://www.ncbi.nlm.nih.gov/sra/47739314 |
| Dmpx_J5_1 | SAMN47739315 | https://www.ncbi.nlm.nih.gov/sra/47739315 |
| Dmpx_J5_2 | SAMN47739316 | https://www.ncbi.nlm.nih.gov/sra/47739316 |
| Dmpx_J5_3 | SAMN47739317 | https://www.ncbi.nlm.nih.gov/sra/47739317 |
| Dmpx_J5_4 | SAMN47739318 | https://www.ncbi.nlm.nih.gov/sra/47739318 |
| Dmpx_J5_5 | SAMN47739319 | https://www.ncbi.nlm.nih.gov/sra/47739319 |
| Dmpx_J6_1 | SAMN47739320 | https://www.ncbi.nlm.nih.gov/sra/47739320 |
| Dmpx_J6_2 | SAMN47739321 | https://www.ncbi.nlm.nih.gov/sra/47739321 |
| Dmpx_J7_1 | SAMN47739322 | https://www.ncbi.nlm.nih.gov/sra/47739322 |
| Dmpx_J7_2 | SAMN47739323 | https://www.ncbi.nlm.nih.gov/sra/47739323 |
| Dmpx_J7_3 | SAMN47739324 | https://www.ncbi.nlm.nih.gov/sra/47739324 |
| Dmpx_J7_4 | SAMN47739325 | https://www.ncbi.nlm.nih.gov/sra/47739325 |
| Dmpx_J7_5 | SAMN47739326 | https://www.ncbi.nlm.nih.gov/sra/47739326 |
| Dmpx_J8_1 | SAMN47739327 | https://www.ncbi.nlm.nih.gov/sra/47739327 |
| Dmpx_J8_2 | SAMN47739328 | https://www.ncbi.nlm.nih.gov/sra/47739328 |
| Dmpx_J8_3 | SAMN47739329 | https://www.ncbi.nlm.nih.gov/sra/47739329 |
| Dmpx_J8_4 | SAMN47739330 | https://www.ncbi.nlm.nih.gov/sra/47739330 |
| Dmpx_J8_5 | SAMN47739331 | https://www.ncbi.nlm.nih.gov/sra/47739331 |
| Dmpx_J9_10 | SAMN47739332 | https://www.ncbi.nlm.nih.gov/sra/47739332 |
| Dmpx_J9_1 | SAMN47739333 | https://www.ncbi.nlm.nih.gov/sra/47739333 |
| Dmpx_J9_2 | SAMN47739334 | https://www.ncbi.nlm.nih.gov/sra/47739334 |
| Dmpx_J9_3 | SAMN47739335 | https://www.ncbi.nlm.nih.gov/sra/47739335 |
| Dmpx_J9_4 | SAMN47739336 | https://www.ncbi.nlm.nih.gov/sra/47739336 |
| Dmpx_J9_5 | SAMN47739337 | https://www.ncbi.nlm.nih.gov/sra/47739337 |
| Dmpx_J9_6 | SAMN47739338 | https://www.ncbi.nlm.nih.gov/sra/47739338 |
| Dmpx_J9_7_r2 | SAMN47739339 | https://www.ncbi.nlm.nih.gov/sra/47739339 |
| Dmpx_J9_8 | SAMN47739340 | https://www.ncbi.nlm.nih.gov/sra/47739340 |
| Dmpx_J9_9 | SAMN47739341 | https://www.ncbi.nlm.nih.gov/sra/47739341 |
| Dmpx_N10_1 | SAMN47739342 | https://www.ncbi.nlm.nih.gov/sra/47739342 |
| Dmpx_N11_1 | SAMN47739343 | https://www.ncbi.nlm.nih.gov/sra/47739343 |
| Dmpx_N11_2 | SAMN47739344 | https://www.ncbi.nlm.nih.gov/sra/47739344 |
| Dmpx_N11_3 | SAMN47739345 | https://www.ncbi.nlm.nih.gov/sra/47739345 |
| Dmpx_N11_4 | SAMN47739346 | https://www.ncbi.nlm.nih.gov/sra/47739346 |
| Dmpx_N11_5 | SAMN47739347 | https://www.ncbi.nlm.nih.gov/sra/47739347 |
| Dmpx_N1_1 | SAMN47739348 | https://www.ncbi.nlm.nih.gov/sra/47739348 |
| Dmpx_N12_1 | SAMN47739349 | https://www.ncbi.nlm.nih.gov/sra/47739349 |
| Dmpx_N12_2 | SAMN47739350 | https://www.ncbi.nlm.nih.gov/sra/47739350 |
| Dmpx_N12_3 | SAMN47739351 | https://www.ncbi.nlm.nih.gov/sra/47739351 |
| Dmpx_N12_4 | SAMN47739352 | https://www.ncbi.nlm.nih.gov/sra/47739352 |
| Dmpx_N12_5 | SAMN47739353 | https://www.ncbi.nlm.nih.gov/sra/47739353 |
| Dmpx_N13_1 | SAMN47739354 | https://www.ncbi.nlm.nih.gov/sra/47739354 |
| Dmpx_N13_2 | SAMN47739355 | https://www.ncbi.nlm.nih.gov/sra/47739355 |
| Dmpx_N13_3 | SAMN47739356 | https://www.ncbi.nlm.nih.gov/sra/47739356 |
| Dmpx_N13_4 | SAMN47739357 | https://www.ncbi.nlm.nih.gov/sra/47739357 |
| Dmpx_N13_5 | SAMN47739358 | https://www.ncbi.nlm.nih.gov/sra/47739358 |
| Dmpx_N14_1 | SAMN47739359 | https://www.ncbi.nlm.nih.gov/sra/47739359 |
| Dmpx_N14_2 | SAMN47739360 | https://www.ncbi.nlm.nih.gov/sra/47739360 |
| Dmpx_N14_3 | SAMN47739361 | https://www.ncbi.nlm.nih.gov/sra/47739361 |
| Dmpx_N14_4 | SAMN47739362 | https://www.ncbi.nlm.nih.gov/sra/47739362 |
| Dmpx_N14_5 | SAMN47739363 | https://www.ncbi.nlm.nih.gov/sra/47739363 |
| Dmpx_N15_1 | SAMN47739364 | https://www.ncbi.nlm.nih.gov/sra/47739364 |
| Dmpx_N15_2 | SAMN47739365 | https://www.ncbi.nlm.nih.gov/sra/47739365 |
| Dmpx_N15_3 | SAMN47739366 | https://www.ncbi.nlm.nih.gov/sra/47739366 |
| Dmpx_N15_4 | SAMN47739367 | https://www.ncbi.nlm.nih.gov/sra/47739367 |
| Dmpx_N15_5 | SAMN47739368 | https://www.ncbi.nlm.nih.gov/sra/47739368 |
| Dmpx_N16_1 | SAMN47739369 | https://www.ncbi.nlm.nih.gov/sra/47739369 |
| Dmpx_N16_2 | SAMN47739370 | https://www.ncbi.nlm.nih.gov/sra/47739370 |
| Dmpx_N16_3 | SAMN47739371 | https://www.ncbi.nlm.nih.gov/sra/47739371 |
| Dmpx_N16_4 | SAMN47739372 | https://www.ncbi.nlm.nih.gov/sra/47739372 |
| Dmpx_N16_5 | SAMN47739373 | https://www.ncbi.nlm.nih.gov/sra/47739373 |
| Dmpx_N17_1 | SAMN47739374 | https://www.ncbi.nlm.nih.gov/sra/47739374 |
| Dmpx_N17_2 | SAMN47739375 | https://www.ncbi.nlm.nih.gov/sra/47739375 |
| Dmpx_N17_3 | SAMN47739376 | https://www.ncbi.nlm.nih.gov/sra/47739376 |
| Dmpx_N17_4 | SAMN47739377 | https://www.ncbi.nlm.nih.gov/sra/47739377 |
| Dmpx_N17_5 | SAMN47739378 | https://www.ncbi.nlm.nih.gov/sra/47739378 |
| Dmpx_N18_1 | SAMN47739379 | https://www.ncbi.nlm.nih.gov/sra/47739379 |
| Dmpx_N19_1 | SAMN47739380 | https://www.ncbi.nlm.nih.gov/sra/47739380 |
| Dmpx_N19_2 | SAMN47739381 | https://www.ncbi.nlm.nih.gov/sra/47739381 |
| Dmpx_N19_3 | SAMN47739382 | https://www.ncbi.nlm.nih.gov/sra/47739382 |
| Dmpx_N19_4 | SAMN47739383 | https://www.ncbi.nlm.nih.gov/sra/47739383 |
| Dmpx_N19_5 | SAMN47739384 | https://www.ncbi.nlm.nih.gov/sra/47739384 |
| Dmpx_N20_1 | SAMN47739385 | https://www.ncbi.nlm.nih.gov/sra/47739385 |
| Dmpx_N20_2 | SAMN47739386 | https://www.ncbi.nlm.nih.gov/sra/47739386 |
| Dmpx_N20_3 | SAMN47739387 | https://www.ncbi.nlm.nih.gov/sra/47739387 |
| Dmpx_N20_4 | SAMN47739388 | https://www.ncbi.nlm.nih.gov/sra/47739388 |
| Dmpx_N20_5 | SAMN47739389 | https://www.ncbi.nlm.nih.gov/sra/47739389 |
| Dmpx_N20_6 | SAMN47739390 | https://www.ncbi.nlm.nih.gov/sra/47739390 |
| Dmpx_N20_7 | SAMN47739391 | https://www.ncbi.nlm.nih.gov/sra/47739391 |
| Dmpx_N20_8 | SAMN47739392 | https://www.ncbi.nlm.nih.gov/sra/47739392 |
| Dmpx_N21_10 | SAMN47739393 | https://www.ncbi.nlm.nih.gov/sra/47739393 |
| Dmpx_N21_1 | SAMN47739394 | https://www.ncbi.nlm.nih.gov/sra/47739394 |
| Dmpx_N21_2 | SAMN47739395 | https://www.ncbi.nlm.nih.gov/sra/47739395 |
| Dmpx_N21_3 | SAMN47739396 | https://www.ncbi.nlm.nih.gov/sra/47739396 |
| Dmpx_N21_4 | SAMN47739397 | https://www.ncbi.nlm.nih.gov/sra/47739397 |
| Dmpx_N21_5 | SAMN47739398 | https://www.ncbi.nlm.nih.gov/sra/47739398 |
| Dmpx_N21_6 | SAMN47739399 | https://www.ncbi.nlm.nih.gov/sra/47739399 |
| Dmpx_N21_7 | SAMN47739400 | https://www.ncbi.nlm.nih.gov/sra/47739400 |
| Dmpx_N21_8 | SAMN47739401 | https://www.ncbi.nlm.nih.gov/sra/47739401 |
| Dmpx_N2_1 | SAMN47739402 | https://www.ncbi.nlm.nih.gov/sra/47739402 |
| Dmpx_N22_10 | SAMN47739403 | https://www.ncbi.nlm.nih.gov/sra/47739403 |
| Dmpx_N22_11 | SAMN47739404 | https://www.ncbi.nlm.nih.gov/sra/47739404 |
| Dmpx_N22_1 | SAMN47739405 | https://www.ncbi.nlm.nih.gov/sra/47739405 |
| Dmpx_N22_2 | SAMN47739406 | https://www.ncbi.nlm.nih.gov/sra/47739406 |
| Dmpx_N22_3 | SAMN47739407 | https://www.ncbi.nlm.nih.gov/sra/47739407 |
| Dmpx_N22_4 | SAMN47739408 | https://www.ncbi.nlm.nih.gov/sra/47739408 |
| Dmpx_N22_5 | SAMN47739409 | https://www.ncbi.nlm.nih.gov/sra/47739409 |
| Dmpx_N22_6 | SAMN47739410 | https://www.ncbi.nlm.nih.gov/sra/47739410 |
| Dmpx_N22_7 | SAMN47739411 | https://www.ncbi.nlm.nih.gov/sra/47739411 |
| Dmpx_N22_8 | SAMN47739412 | https://www.ncbi.nlm.nih.gov/sra/47739412 |
| Dmpx_N22_9 | SAMN47739413 | https://www.ncbi.nlm.nih.gov/sra/47739413 |
| Dmpx_N23_10 | SAMN47739414 | https://www.ncbi.nlm.nih.gov/sra/47739414 |
| Dmpx_N23_1 | SAMN47739415 | https://www.ncbi.nlm.nih.gov/sra/47739415 |
| Dmpx_N23_2 | SAMN47739416 | https://www.ncbi.nlm.nih.gov/sra/47739416 |
| Dmpx_N23_3 | SAMN47739417 | https://www.ncbi.nlm.nih.gov/sra/47739417 |
| Dmpx_N23_4 | SAMN47739418 | https://www.ncbi.nlm.nih.gov/sra/47739418 |
| Dmpx_N23_5 | SAMN47739419 | https://www.ncbi.nlm.nih.gov/sra/47739419 |
| Dmpx_N23_6 | SAMN47739420 | https://www.ncbi.nlm.nih.gov/sra/47739420 |
| Dmpx_N23_7 | SAMN47739421 | https://www.ncbi.nlm.nih.gov/sra/47739421 |
| Dmpx_N23_8 | SAMN47739422 | https://www.ncbi.nlm.nih.gov/sra/47739422 |
| Dmpx_N23_9 | SAMN47739423 | https://www.ncbi.nlm.nih.gov/sra/47739423 |
| Dmpx_N24_10 | SAMN47739424 | https://www.ncbi.nlm.nih.gov/sra/47739424 |
| Dmpx_N24_11 | SAMN47739425 | https://www.ncbi.nlm.nih.gov/sra/47739425 |
| Dmpx_N24_12 | SAMN47739426 | https://www.ncbi.nlm.nih.gov/sra/47739426 |
| Dmpx_N24_1 | SAMN47739427 | https://www.ncbi.nlm.nih.gov/sra/47739427 |
| Dmpx_N24_2 | SAMN47739428 | https://www.ncbi.nlm.nih.gov/sra/47739428 |
| Dmpx_N24_3 | SAMN47739429 | https://www.ncbi.nlm.nih.gov/sra/47739429 |
| Dmpx_N24_4 | SAMN47739430 | https://www.ncbi.nlm.nih.gov/sra/47739430 |
| Dmpx_N24_5 | SAMN47739431 | https://www.ncbi.nlm.nih.gov/sra/47739431 |
| Dmpx_N24_6 | SAMN47739432 | https://www.ncbi.nlm.nih.gov/sra/47739432 |
| Dmpx_N24_7 | SAMN47739433 | https://www.ncbi.nlm.nih.gov/sra/47739433 |
| Dmpx_N24_8 | SAMN47739434 | https://www.ncbi.nlm.nih.gov/sra/47739434 |
| Dmpx_N24_9 | SAMN47739435 | https://www.ncbi.nlm.nih.gov/sra/47739435 |
| Dmpx_N25_1 | SAMN47739436 | https://www.ncbi.nlm.nih.gov/sra/47739436 |
| Dmpx_N25_2 | SAMN47739437 | https://www.ncbi.nlm.nih.gov/sra/47739437 |
| Dmpx_N25_3 | SAMN47739438 | https://www.ncbi.nlm.nih.gov/sra/47739438 |
| Dmpx_N25_4 | SAMN47739439 | https://www.ncbi.nlm.nih.gov/sra/47739439 |
| Dmpx_N25_5 | SAMN47739440 | https://www.ncbi.nlm.nih.gov/sra/47739440 |
| Dmpx_N3_1 | SAMN47739441 | https://www.ncbi.nlm.nih.gov/sra/47739441 |
| Dmpx_N3_2 | SAMN47739442 | https://www.ncbi.nlm.nih.gov/sra/47739442 |
| Dmpx_N3_3 | SAMN47739443 | https://www.ncbi.nlm.nih.gov/sra/47739443 |
| Dmpx_N3_4 | SAMN47739444 | https://www.ncbi.nlm.nih.gov/sra/47739444 |
| Dmpx_N3_5 | SAMN47739445 | https://www.ncbi.nlm.nih.gov/sra/47739445 |
| Dmpx_N4_1 | SAMN47739446 | https://www.ncbi.nlm.nih.gov/sra/47739446 |
| Dmpx_N4_2 | SAMN47739447 | https://www.ncbi.nlm.nih.gov/sra/47739447 |
| Dmpx_N4_3 | SAMN47739448 | https://www.ncbi.nlm.nih.gov/sra/47739448 |
| Dmpx_N4_4 | SAMN47739449 | https://www.ncbi.nlm.nih.gov/sra/47739449 |
| Dmpx_N4_5 | SAMN47739450 | https://www.ncbi.nlm.nih.gov/sra/47739450 |
| Dmpx_N5_1 | SAMN47739451 | https://www.ncbi.nlm.nih.gov/sra/47739451 |
| Dmpx_N5_2 | SAMN47739452 | https://www.ncbi.nlm.nih.gov/sra/47739452 |
| Dmpx_N5_3 | SAMN47739453 | https://www.ncbi.nlm.nih.gov/sra/47739453 |
| Dmpx_N5_4 | SAMN47739454 | https://www.ncbi.nlm.nih.gov/sra/47739454 |
| Dmpx_N5_5 | SAMN47739455 | https://www.ncbi.nlm.nih.gov/sra/47739455 |
| Dmpx_N6_1 | SAMN47739456 | https://www.ncbi.nlm.nih.gov/sra/47739456 |
| Dmpx_N6_2 | SAMN47739457 | https://www.ncbi.nlm.nih.gov/sra/47739457 |
| Dmpx_N6_3 | SAMN47739458 | https://www.ncbi.nlm.nih.gov/sra/47739458 |
| Dmpx_N6_4 | SAMN47739459 | https://www.ncbi.nlm.nih.gov/sra/47739459 |
| Dmpx_N6_5 | SAMN47739460 | https://www.ncbi.nlm.nih.gov/sra/47739460 |
| Dmpx_N7_1 | SAMN47739461 | https://www.ncbi.nlm.nih.gov/sra/47739461 |
| Dmpx_N8_1 | SAMN47739462 | https://www.ncbi.nlm.nih.gov/sra/47739462 |
| Dmpx_N8_2 | SAMN47739463 | https://www.ncbi.nlm.nih.gov/sra/47739463 |
| Dmpx_N8_3 | SAMN47739464 | https://www.ncbi.nlm.nih.gov/sra/47739464 |
| Dmpx_N8_4 | SAMN47739465 | https://www.ncbi.nlm.nih.gov/sra/47739465 |
| Dmpx_N8_5 | SAMN47739466 | https://www.ncbi.nlm.nih.gov/sra/47739466 |
| Dmpx_N9_1 | SAMN47739467 | https://www.ncbi.nlm.nih.gov/sra/47739467 |
| Dmpx_N9_2 | SAMN47739468 | https://www.ncbi.nlm.nih.gov/sra/47739468 |
| Dmpx_N9_3 | SAMN47739469 | https://www.ncbi.nlm.nih.gov/sra/47739469 |
| Dmpx_N9_4 | SAMN47739470 | https://www.ncbi.nlm.nih.gov/sra/47739470 |
| Dmpx_N9_5 | SAMN47739471 | https://www.ncbi.nlm.nih.gov/sra/47739471 |
| Dmpx_S10_1 | SAMN47739472 | https://www.ncbi.nlm.nih.gov/sra/47739472 |
| Dmpx_S10_2 | SAMN47739473 | https://www.ncbi.nlm.nih.gov/sra/47739473 |
| Dmpx_S10_3 | SAMN47739474 | https://www.ncbi.nlm.nih.gov/sra/47739474 |
| Dmpx_S10_4 | SAMN47739475 | https://www.ncbi.nlm.nih.gov/sra/47739475 |
| Dmpx_S10_5 | SAMN47739476 | https://www.ncbi.nlm.nih.gov/sra/47739476 |
| Dmpx_S11_1 | SAMN47739477 | https://www.ncbi.nlm.nih.gov/sra/47739477 |
| Dmpx_S1_1 | SAMN47739478 | https://www.ncbi.nlm.nih.gov/sra/47739478 |
| Dmpx_S12_1 | SAMN47739479 | https://www.ncbi.nlm.nih.gov/sra/47739479 |
| Dmpx_S12_2 | SAMN47739480 | https://www.ncbi.nlm.nih.gov/sra/47739480 |
| Dmpx_S12_3 | SAMN47739481 | https://www.ncbi.nlm.nih.gov/sra/47739481 |
| Dmpx_S12_4 | SAMN47739482 | https://www.ncbi.nlm.nih.gov/sra/47739482 |
| Dmpx_S12_5 | SAMN47739483 | https://www.ncbi.nlm.nih.gov/sra/47739483 |
| Dmpx_S1_2 | SAMN47739484 | https://www.ncbi.nlm.nih.gov/sra/47739484 |
| Dmpx_S13_1 | SAMN47739485 | https://www.ncbi.nlm.nih.gov/sra/47739485 |
| Dmpx_S13_2 | SAMN47739486 | https://www.ncbi.nlm.nih.gov/sra/47739486 |
| Dmpx_S13_3 | SAMN47739487 | https://www.ncbi.nlm.nih.gov/sra/47739487 |
| Dmpx_S13_4 | SAMN47739488 | https://www.ncbi.nlm.nih.gov/sra/47739488 |
| Dmpx_S13_5 | SAMN47739489 | https://www.ncbi.nlm.nih.gov/sra/47739489 |
| Dmpx_S1_3 | SAMN47739490 | https://www.ncbi.nlm.nih.gov/sra/47739490 |
| Dmpx_S1_4 | SAMN47739491 | https://www.ncbi.nlm.nih.gov/sra/47739491 |
| Dmpx_S1_5 | SAMN47739492 | https://www.ncbi.nlm.nih.gov/sra/47739492 |
| Dmpx_S2_1 | SAMN47739493 | https://www.ncbi.nlm.nih.gov/sra/47739493 |
| Dmpx_S2_2 | SAMN47739494 | https://www.ncbi.nlm.nih.gov/sra/47739494 |
| Dmpx_S2_3 | SAMN47739495 | https://www.ncbi.nlm.nih.gov/sra/47739495 |
| Dmpx_S2_4 | SAMN47739496 | https://www.ncbi.nlm.nih.gov/sra/47739496 |
| Dmpx_S2_5 | SAMN47739497 | https://www.ncbi.nlm.nih.gov/sra/47739497 |
| Dmpx_S3_10 | SAMN47739498 | https://www.ncbi.nlm.nih.gov/sra/47739498 |
| Dmpx_S3_1 | SAMN47739499 | https://www.ncbi.nlm.nih.gov/sra/47739499 |
| Dmpx_S3_2 | SAMN47739500 | https://www.ncbi.nlm.nih.gov/sra/47739500 |
| Dmpx_S3_3 | SAMN47739501 | https://www.ncbi.nlm.nih.gov/sra/47739501 |
| Dmpx_S3_4 | SAMN47739502 | https://www.ncbi.nlm.nih.gov/sra/47739502 |
| Dmpx_S3_5 | SAMN47739503 | https://www.ncbi.nlm.nih.gov/sra/47739503 |
| Dmpx_S3_6 | SAMN47739504 | https://www.ncbi.nlm.nih.gov/sra/47739504 |
| Dmpx_S3_7 | SAMN47739505 | https://www.ncbi.nlm.nih.gov/sra/47739505 |
| Dmpx_S3_8 | SAMN47739506 | https://www.ncbi.nlm.nih.gov/sra/47739506 |
| Dmpx_S3_9 | SAMN47739507 | https://www.ncbi.nlm.nih.gov/sra/47739507 |
| Dmpx_S4_1 | SAMN47739508 | https://www.ncbi.nlm.nih.gov/sra/47739508 |
| Dmpx_S4_2 | SAMN47739509 | https://www.ncbi.nlm.nih.gov/sra/47739509 |
| Dmpx_S4_3 | SAMN47739510 | https://www.ncbi.nlm.nih.gov/sra/47739510 |
| Dmpx_S4_4 | SAMN47739511 | https://www.ncbi.nlm.nih.gov/sra/47739511 |
| Dmpx_S4_5 | SAMN47739512 | https://www.ncbi.nlm.nih.gov/sra/47739512 |
| Dmpx_S5_10 | SAMN47739513 | https://www.ncbi.nlm.nih.gov/sra/47739513 |
| Dmpx_S5_1 | SAMN47739514 | https://www.ncbi.nlm.nih.gov/sra/47739514 |
| Dmpx_S5_2 | SAMN47739515 | https://www.ncbi.nlm.nih.gov/sra/47739515 |
| Dmpx_S5_3 | SAMN47739516 | https://www.ncbi.nlm.nih.gov/sra/47739516 |
| Dmpx_S5_4 | SAMN47739517 | https://www.ncbi.nlm.nih.gov/sra/47739517 |
| Dmpx_S5_5 | SAMN47739518 | https://www.ncbi.nlm.nih.gov/sra/47739518 |
| Dmpx_S5_6 | SAMN47739519 | https://www.ncbi.nlm.nih.gov/sra/47739519 |
| Dmpx_S5_7 | SAMN47739520 | https://www.ncbi.nlm.nih.gov/sra/47739520 |
| Dmpx_S5_8 | SAMN47739521 | https://www.ncbi.nlm.nih.gov/sra/47739521 |
| Dmpx_S5_9 | SAMN47739522 | https://www.ncbi.nlm.nih.gov/sra/47739522 |
| Dmpx_S6_10 | SAMN47739523 | https://www.ncbi.nlm.nih.gov/sra/47739523 |
| Dmpx_S6_11 | SAMN47739524 | https://www.ncbi.nlm.nih.gov/sra/47739524 |
| Dmpx_S6_12 | SAMN47739525 | https://www.ncbi.nlm.nih.gov/sra/47739525 |
| Dmpx_S6_13 | SAMN47739526 | https://www.ncbi.nlm.nih.gov/sra/47739526 |
| Dmpx_S6_14 | SAMN47739527 | https://www.ncbi.nlm.nih.gov/sra/47739527 |
| Dmpx_S6_15 | SAMN47739528 | https://www.ncbi.nlm.nih.gov/sra/47739528 |
| Dmpx_S6_6 | SAMN47739529 | https://www.ncbi.nlm.nih.gov/sra/47739529 |
| Dmpx_S6_7 | SAMN47739530 | https://www.ncbi.nlm.nih.gov/sra/47739530 |
| Dmpx_S6_8 | SAMN47739531 | https://www.ncbi.nlm.nih.gov/sra/47739531 |
| Dmpx_S6_9 | SAMN47739532 | https://www.ncbi.nlm.nih.gov/sra/47739532 |
| Dmpx_S7_1 | SAMN47739533 | https://www.ncbi.nlm.nih.gov/sra/47739533 |
| Dmpx_S7_2 | SAMN47739534 | https://www.ncbi.nlm.nih.gov/sra/47739534 |
| Dmpx_S7_3 | SAMN47739535 | https://www.ncbi.nlm.nih.gov/sra/47739535 |
| Dmpx_S7_4 | SAMN47739536 | https://www.ncbi.nlm.nih.gov/sra/47739536 |
| Dmpx_S7_5 | SAMN47739537 | https://www.ncbi.nlm.nih.gov/sra/47739537 |
| Dmpx_S8_1 | SAMN47739538 | https://www.ncbi.nlm.nih.gov/sra/47739538 |
| Dmpx_S8_2 | SAMN47739539 | https://www.ncbi.nlm.nih.gov/sra/47739539 |
| Dmpx_S8_3 | SAMN47739540 | https://www.ncbi.nlm.nih.gov/sra/47739540 |
| Dmpx_S8_4 | SAMN47739541 | https://www.ncbi.nlm.nih.gov/sra/47739541 |
| Dmpx_S8_5 | SAMN47739542 | https://www.ncbi.nlm.nih.gov/sra/47739542 |
| Dmpx_S9_1 | SAMN47739543 | https://www.ncbi.nlm.nih.gov/sra/47739543 |
| Dmpx_S9_2 | SAMN47739544 | https://www.ncbi.nlm.nih.gov/sra/47739544 |
| Dmpx_S9_3 | SAMN47739545 | https://www.ncbi.nlm.nih.gov/sra/47739545 |
| Dmpx_S9_4 | SAMN47739546 | https://www.ncbi.nlm.nih.gov/sra/47739546 |
| Dmpx_S9_5 | SAMN47739547 | https://www.ncbi.nlm.nih.gov/sra/47739547 |
| Dmpx_XSP_1 | SAMN47739548 | https://www.ncbi.nlm.nih.gov/sra/47739548 |
| Dmpx_XSP_2 | SAMN47739549 | https://www.ncbi.nlm.nih.gov/sra/47739549 |
| Dmpx_XSP_3 | SAMN47739550 | https://www.ncbi.nlm.nih.gov/sra/47739550 |
| Dmpx_XSP_4 | SAMN47739551 | https://www.ncbi.nlm.nih.gov/sra/47739551 |
| Dmpx_XSP_5 | SAMN47739552 | https://www.ncbi.nlm.nih.gov/sra/47739552 |
